# Increased intracellular persulfide levels attenuate HlyU-mediated hemolysin transcriptional activation in *Vibrio cholerae*

**DOI:** 10.1101/2023.03.13.532278

**Authors:** Cristian M. Pis Diez, Giuliano T. Antelo, Triana N. Dalia, Ankur B. Dalia, David P. Giedroc, Daiana A. Capdevila

**Author notes:** Correspondence: Prof. Daiana A. Capdevila, Fundación Instituto Leloir, Instituto de Investigaciones Bioquímicas de Buenos Aires (IIBBA-CONICET), C1405BWE Ciudad Autónoma de, Buenos Aires, Argentina. Prof. David P. Giedroc, Indiana University, Department of Chemistry, 800 E. Kirkwood Avenue, Bloomington, IN 47405-7102 USA.

## Abstract

The vertebrate host’s immune system and resident commensal bacteria deploy a range of highly reactive small molecules that provide a barrier against infections by microbial pathogens. Gut pathogens, such as *Vibrio cholerae*, sense and respond to these stressors by modulating the expression of exotoxins that are crucial for colonization. Here, we employ mass-spectrometry-based profiling, metabolomics, expression assays and biophysical approaches to show that transcriptional activation of the hemolysin gene *hlyA* in *V. cholerae* is regulated by intracellular reactive sulfur species (RSS), specifically sulfane sulfur. We first present a comprehensive sequence similarity network analysis of the arsenic repressor (ArsR) superfamily of transcriptional regulators where RSS and reactive oxygen species (ROS) sensors segregate into distinct clusters. We show that HlyU, transcriptional activator of *hlyA* in *V. cholerae*, belongs to the RSS-sensing cluster and readily reacts with organic persulfides, showing no reactivity and remaining DNA-bound following treatment with various ROS in vitro, including H_2_O_2_. Surprisingly, in *V. cholerae* cell cultures, both sulfide and peroxide treatment downregulate HlyU-dependent transcriptional activation of *hlyA*. However, RSS metabolite profiling shows that both sulfide and peroxide treatment raise the endogenous inorganic sulfide and disulfide levels to a similar extent, accounting for this crosstalk, and confirming that *V. cholerae* attenuates HlyU-mediated activation of *hlyA* in a specific response to intracellular RSS. These findings provide new evidence that gut pathogens may harness RSS-sensing as an evolutionary adaptation that allows them to overcome the gut inflammatory response by modulating the expression of exotoxins.

## INTRODUCTION

Many bacterial pathogens secrete diverse protein toxins to disrupt host defense systems, which are expressed with precise spatiotemporal regulation, since untimely toxin secretion can be detrimental to the invading pathogens (1, 2). Such is the case of *Vibrio cholerae*, the major causative agent of the severe diarrheal disease, cholera. In this organism, the expression of various enteric exotoxins is under control of distinct transcriptional regulators that trigger their expression upon attachment to the small intestine epithelium surface, enabling efficient colonization (3). Cholera toxin (CT), the major virulence factor responsible for cholera pathogenesis, and other accessory toxin and virulence factors (*e.g.*, Ace, Zot, TCP) are primarily regulated by the transcriptional activator ToxT (4). Extensive *in vitro* and *ex vivo* studies have revealed that ToxT activation is prevented by components of the bile found in the gut, which bind directly to ToxT or to other proteins of the regulatory network, allowing the expression of ToxT-activated virulence factors only upon reaching the epithelium surface (5–8). Beyond ToxT activated genes, pathogenic strains of *V. cholerae* produce several additional accessory toxins (2) such as the extracellular pore-forming toxin hemolysin (HlyA) (9), which is implicated in pathogenesis, particularly, in those strains that lack CT (10). HlyA has been reported to be activated by HlyU and repressed by the quorum sensing regulator HapR and the iron uptake repressor Fur (11). While HapR and Fur link quorum sensing (12) and cellular iron status (13) to virulence gene regulation, which is likely advantageous in the human host, the signals that modulate HlyU-dependent activation of HlyA in *V. cholerae* and other exotoxins in other *Vibrio species* remain unresolved (14–24). Here, we present a biochemical and functional characterization of HlyU-mediated responses towards microenvironmental signals thought to be present in the gut that may impact hemolysin expression of *V*. *cholerae* and other exotoxins in pathogenic *Vibrio* species (18–20, 25).

To cause cholera, *V. cholerae* must effectively colonize the small intestine (SI), overcoming many stress conditions (3). First, acid tolerance factors along with biofilm formation serves as adaptation to the low pH of the human stomach (26, 27). Upon reaching the SI, efflux pumps and changes in motility allow *Vibrio* to repel liver-derived bile and antimicrobial peptides in the intestinal lumen (28, 29). Then, mucolytic agents and proteases help penetrate a highly viscous mucus layer that acts as physical barrier of the intestinal epithelium for bacterial colonization (30). Finally, once reaching the epithelium *V. cholerae* produces toxins while facing the inflammatory response of the host tissue (7, 31). More recent work shows that gut pathogens must also overcome hydrogen sulfide-stress imposed by the host or gut microbiota (32), which involves an increase in bile acid (taurocholic acid)-derived hydrogen sulfide (H_2_S) and potentially more oxidized reactive sulfur species (RSS) (33, 34). Although gut pathogen adaptation to such stressors remains understudied, the protein machinery charged with H_2_S/RSS remediation and, ultimately, efflux has been described for many bacterial pathogens and free-living bacteria (35–40). It is now clear that beneficial levels of H_2_S and low molecular weight thiol persulfides can protect bacteria from oxidative stress that arises from inflammatory responses (41–43), as well as antibiotics (44–46). Recently, it has been shown that *V. cholerae* produces endogenous H_2_S, which decreases its susceptibility to H_2_O_2_ in both *in vitro* and *in vivo* adult mice models (47). Thus, beyond exposure to exogenous H_2_S (32), intracellular H_2_S/RSS levels in *V. cholerae* also depend on the synthesis of endogenous H_2_S from cysteine metabolism. We speculate that exogenous and endogenous levels of H_2_S may serve as additional microenvironmental cues that impact SI colonization dynamics and toxin gene expression.

H_2_S is as an important signaling molecule for gut bacteria. However, little is known about the intracellular concentration of the components of the RSS pool, something particularly true for *V. cholerae*. This encompasses organic and inorganic molecules containing sulfur in higher oxidation states than H_2_S, many of which contain sulfur-bonded or “sulfane” sulfur (34). These species are both responsible for the beneficial biological properties of H_2_S, as well as its toxicity at least in part, as RSS can effectively modify catalytic residues in proteins, often negatively impacting their activity (37, 48, 49). Signaling by RSS is achieved predominantly via a post-translational modification of cysteine residues in proteins, often referred to as *S*-sulfuration, persulfidation or *S*-sulfhydration (49). Thus, the speciation of “sulfane” sulfur inside cells, meaning the cellular concentrations of all organic, inorganic and protein species, must be controlled. Bacteria maintain H_2_S/RSS homeostasis by expressing persulfide-sensing transcriptional regulators which regulons generally encode for a subset of common downstream H_2_S detoxification genes (50, 51). The mechanisms that allow H_2_S/RSS homeostasis in bacteria have been described for several human pathogens (34, 37, 48, 52), however little is known about how *V. cholerae* responds to increasing H_2_S/RSS and how these reactive species affect pathogen metabolism and, ultimately, gut colonization.

The described persulfide-sensing transcriptional regulators in bacteria belong to three structurally unrelated protein families, namely the arsenic repressor (ArsR), copper-sensitive operon repressor (CsoR) and Fis superfamilies. They all harness dithiol chemistry to form either disulfide or polysulfide bridges between reactive cysteine (Cys) residues that allow for the transcription of sulfide metabolism genes, either by transcriptional derepression or by RNA polymerase recruitment and transcriptional activation (48, 50, 51, 53). Beyond *bona fide* persulfide-sensing transcriptional regulators that control the expression of at least one sulfide metabolism gene, other dithiol-harboring transcriptional regulators have been reported to react and elicit transcriptional responses in the presence of persulfides and/or polysulfides (54–57). To what extent these sensors are truly specific to persulfides and can be distinguished from other redox sensors that readily react with H_2_O_2_ *in vitro* remains a matter of debate despite the detailed structural information available (50, 51). Nevertheless, it is interesting to evaluate persulfide-sensing as an evolutionary advantage for human pathogens as these species are prevalent and biosynthesized in certain tissues by the host or host-resident microbiota, notably in the gut (58).

To date, no persulfide-sensing transcriptional regulator has been identified in *Vibrio* strains, thus we aimed to predict putative regulators that would respond to these species by sequence homology of ArsR, CsoR and Fis family proteins encoded in the *V. cholerae* genome. While *V. cholerae* strains do not generally encode for CsoR (51) or FisR family proteins (59), they encode at least two ArsR family proteins, one of which is HlyU and the other one annotated as BigR (biofilm repressor) in some strains (60). Both proteins conserve the Cys pair that functions as the sensing site in *Rhodobacter capsulatus* SqrR and *Acinetobacter baumanii* BigR, two extensively characterized persulfide-sensing transcriptional regulators from the ArsR family (35, 48, 50). These RSS-sensing repressors react with both organic and inorganic persulfides, but not with disulfides or peroxides, leading to the formation of a polysulfide bridge between the cysteines (50), and are readily distinguished from other ArsR-family members that harbor H_2_O_2_-sensing proximal cysteine site (61). While *Vc*BigR and other ArsR family proteins in *V. cholera* remain completely uncharacterized, the structure of *Vc*HlyU is known and the role of cysteine oxidation in inhibiting DNA operator binding has been previously reported (62). However, it remains unclear whether HlyU senses ROS levels in its local environment since Cys sulfenylation may not be prevalent in the gastrointestinal tract due to the microaerobic or anaerobic conditions. In this context, the sequence similarities between HlyU and previously characterized *bona fide* persulfide sensors strongly motivate experiments capable of defining the chemical specificity of this transcriptional regulator and, ultimately, understand the molecular basis of how HlyU oxidative modifications prevent the initiation of the virulence cascade in *Vibrio* spp.

In this study, we first use a comprehensive sequence similarity network (SSN) analysis of >150,000 sequences to identify distinct clusters of RSS-sensitive and ROS-sensitive regulators, which reveals that HlyU is most closely related to previously characterized persulfide sensors (50). Next, we investigated the reactivity of HlyU using *in vitro* mass spectrometry and fluorescence anisotropy, which show that HlyU reacts with organic persulfides to form a tetrasulfide bridge between its two cysteines, abrogating DNA-binding. In striking contrast, disulfides and peroxides do not react with HlyU nor do they impact DNA binding. We found that the exogenous treatment of *V. cholerae* cells with either sulfide (Na_2_S) or hydrogen peroxide (H_2_O_2_) attenuates HlyU-dependent activation of *hlyA*, downregulating *hlyA* transcription likely through transcriptional silencing by the nucleoprotein H-NS (63). Quantitative RSS metabolite profiling experiments reveal that both treatments result in an increase in the levels of inorganic persulfides thus reconciling the *in vitro* and *in vivo* results. Together, these findings suggest that persulfides function as the cognate regulator of HlyU-regulated gene expression, thus uncovering a new role for RSS sensing in exotoxin expression in a major enteric pathogen.

## RESULTS

### Sequence similarity network (SSN) analysis of the ArsR superfamily suggests that HlyU is an RSS-sensing transcriptional regulator

ArsR superfamily proteins are compact homodimeric “winged helical” DNA binding proteins characterized by a core α1-α2-α3-α4-β1-β2-α5 secondary structure; some members contain extensions on either or both N- and C-terminal sides of this motif, and if α-helical are denoted the α0 and α6 helices, respectively, to facilitate sequence comparisons (64, 65). The helix-turn-helix (HTH) motif that mediates DNA-binding is the α3-α4 segment that engages successive DNA major grooves, with a degenerate tetrapeptide sequence in α4 (Figure S1A) that enforces specificity for a particular DNA operator sequence (66). The β1-β2 wing extends from the periphery of the dimer and may mediate interactions with the immediately adjacent minor grooves. Early work on ArsR family proteins suggested that the >3000 distinct members of this family were mostly metal ion or metalloid-specific regulators (67–69). However, it has become increasingly clear that this is not the case (65, 70) as many recently described regulators have been shown to respond to reactive inorganic species (35, 40, 61) or may lack inducer binding sites altogether (71–73). To understand the sequence relationships between these proteins and evaluate to what extent the differences in inducer recognition sites are captured in a large superfamily with low pairwise sequence conservation, we performed a sequence similarity network (SSN) analysis (74) of 168,163 unique entries from the Pfam PF01022 and Interpro IPR001845 datasets (13,879 sequences that are <50% identical over 80% of the sequence; UNIREF50 clusters). Functionally characterized members in each cluster suggest that individual clusters may represent groups of proteins that share the same chemical inducer (Figure 1A, Figure S1A and Supplementary Table S2) which is further supported by the conservation of inducer recognition residues in well-described distinct structural motifs within each SSN cluster (Figure 1B) (65).

**Figure 1.**
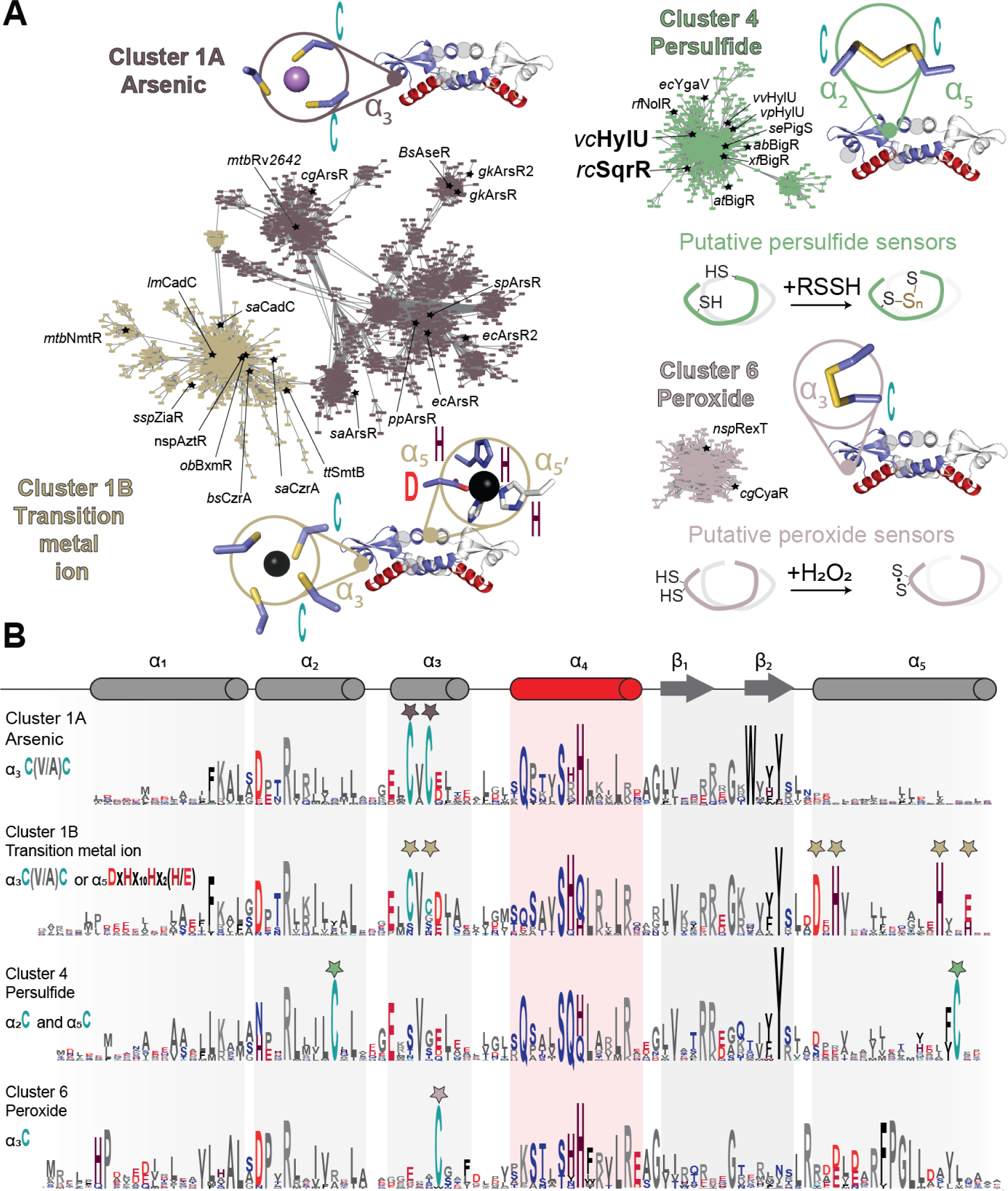
Sequence similarity network (SSN) analysis of the ArsR superfamily of bacterial repressors. **(A)** Results of an SSN clustering analysis of 168,163 unique sequences belonging to the Pfam PF01022 and Interpro IPR001845 Pfam using genomic enzymology tools and visualized using Cytoscape. Clusters functionally annotated as arsenic (As), nickel/cobalt (Ni/Co), persulfide (RSSH) or hydrogen peroxide (H_2_O_2_) sensors are presented here, and all the determined main clusters are presented in Fig. S1. All clusters are designated by a number and ranked according to the number of unique sequences (Table S2), color-coded, and, if known. Each node corresponds to sequences that are 50% identical over 80% of the sequence, using an alignment score of 22 (see Materials and Methods). Functionally characterized members in each are indicated with species name and trivial name. **(B)** Sequence logo representations of sequence conservation defined by the indicated cluster of sequences derived from panel A. The residues that coordinate metals/metalloid ions in ArsR are marked by stars and the residues that undergo redox chemistry are marked with arrows. WebLogos other clusters are also provided (Figure S1B).

The largest SSN cluster that appears at this level of sequence segregation (see Methods) is cluster 1, which can be divided into two large subclusters, denoted here as 1A and 1B. These sequences share a highly conserved CXC motif in the third helix (α3) (Figure 1A, *dark brown*), and are distinguished from one another by the absence (subcluster 1A) or presence (subcluster 1B) of an additional inducer recognition site known to bind transition metals (Figure 1A; *gold*) (65). Cluster 1, in fact, contains the vast majority of described metalloregulators to date, that sense either biological transition metals, *e.g*., Cu(I), Zn(II), Ni(II) or Co(II) and heavy metal xenobiotics Cd(II) and Pb(II) (subcluster 1B), or trivalent As(III)/Sb(III) (subcluster 1A). The exceptions are the sequences compiled in SSN cluster 21, representative of the Ni(II) sensor SrnR (73) and those in SSN cluster 22, representative of the Cd(II) sensor CmtR, with metal coordinating ligands derived from the DNA binding helix α4 (Figure S1B) (75, 76). Subcluster 1B contains all historically characterized, canonical As(III) sensors harboring a C-(V/A)-C motif that coordinates As(III), while cluster 5 includes many more recently described atypical As(III) sensors that feature trigonal Cys coordination with all metal ligands derived from α5 (Figure S1B) (77).

The remainder of the SSN clusters represent proteins that are not obviously metal ion or metalloid-sensing regulators. SSN clusters 2 and 3 (Figure S1B) remain largely uncharacterized and contain members involved in some way in the hypoxic response in *M. tuberculosis* (Rv2034 and Rv0081, respectively, Supplementary Table S2) (72, 78). SSN cluster 4 encompasses all known RSS or persulfide sensors characterized by a pair of cysteines in the α2 and α5 helices which form an intraprotomer polysulfide bridge when presented with sulfane sulfur donors (48, 50). As expected from a previous SSN (61), cluster 4 sequences are readily distinguished from those in SSN cluster 6, which contains a recently characterized dithiol protein, RexT, that reacts with H_2_O_2_ via two proximal Cys residues in α3. Overall, our SSN analysis suggests that ArsR proteins that lack a metal binding site can be readily distinguished and may indeed harness certain degree of chemical specificity of distinct dithiol sites. Moreover, the two ArsR protein encoded by *V. cholerae*, HlyU (locus tag VC_0678) and BigR (VC_0642) are members of SSN cluster 4, thus suggesting they may respond primarily to inorganic and organic persulfides, and not to hypoxia as described for proteins in SSN clusters 2 and 3, nor H_2_O_2_ as it has been described for SSN cluster 6.

These functional assignments are further supported by a genome neighborhood analysis, based on the premise that in bacteria, regulatory and functional genes dedicated to a particular task tend to form gene clusters in the chromosome. While the genes encoding the metalloregulators in SSN clusters 1A, 21 and 22 are generally nearby one or more genes encoding a metal ion transporter, the neighboring genes of As(III)-dependent repressors (clusters 1B and 5) encode for arsenate-transferring proteins and organoarsenic transporters (Figure S2). Similarly, known persulfide-sensing cluster 4 regulators genomically co-localize with genes encoding sulfurtransferases or rhodaneses and other sulfur-oxidizing persulfide-reducing enzymes and inorganic sulfur transporters (36, 37, 48). In contrast, genes encoding peroxide-sensing SSN cluster 6 regulators are generally nearby genes encoding NADH oxidoreductases. A more comprehensive analysis of the regulons of SSN cluster 4 persulfide-sensing repressors suggests that exotoxin expression in *Vibrio* ssp., and biofilm production and antibiotic biosynthesis in others is linked in some way to RSS-sensing in cells, a remarkably diverse collection of adaptive responses that are likely tuned to bacterial lifestyle and environmental needs. In this context, we characterize *V. cholerae* HlyU, the positive regulator of hemolysin production. The inducer-selectivity of HlyU remains undefined, although previous work implicates reactive oxygen species in this role, in striking contrast to the implications of SSN analysis which places HlyU in SSN cluster 4 (22).

### HlyU reacts exclusively with persulfides to form a tetrasulfide bridge that leads to reversible DNA dissociation

To evaluate biochemically if sulfide signaling through persulfides impacts exotoxin expression in *V. cholerae* through a HlyU-dependent mechanism, we determined the ability of HlyU to distinguish between persulfides and other non-sulfur containing oxidants. We exploited a mass spectrometry (MS)-based, anaerobic assay (50, 79) to determine the reactivity of HlyU toward redox-active small molecules in a time-resolved manner. In this assay, we employ quantitative capping by excess iodoacetamide (IAM) in the absence of denaturing agents, as confirmation that the protein is fully reduced. Excess IAM is removed immediately after quenching to prevent undesired chemistry of the capping agent (80) (Fig. 2A). Consistent with expectations from the placement of HlyU among other persulfide sensors in SSN cluster 4 (50), we observed no change in the mass spectrum upon IAM capping following a one-hour incubation with a 20-fold molar excess of H_2_O_2_, and glutathione disulfide GSSG (Figs. 2A, 2C and S3). In contrast, reduced HlyU readily reacts with inorganic (Na_2_S_4_) and organic (glutathione persulfide, GSSH; cysteine persulfide, CSSH; homocysteine persulfide, hCSSH) persulfides, shifting the mass distribution to a +62-Da species, consistent with an intramolecular (intraprotomer) tetrasulfide crosslink between the conserved Cys38 and Cys101 residues (Fig. 2A, 2B and S3). Interestingly, HlyU tetrasulfide differs from previously reported experiments carried out with SqrR and *Ab*BigR, in that the tetrasulfide can be partially capped with IAM. This suggests that the tetrasulfide is in equilibrium with an “open” hydropersulfide or hydropolysulfide form. Moreover, to rule out the formation of the sulfenic acid on Cys38 previously captured by crystallography in air in absence of a reducing agent (62), we performed overnight aerobic reactions with IAM or dimedone. These experiments confirm that both Cys remain reduced when treated with H_2_O_2_ and are capable only of forming a tetrasulfide bridge when treated with sulfane sulfur donors (Fig. 2B-C).

**Figure 2.**
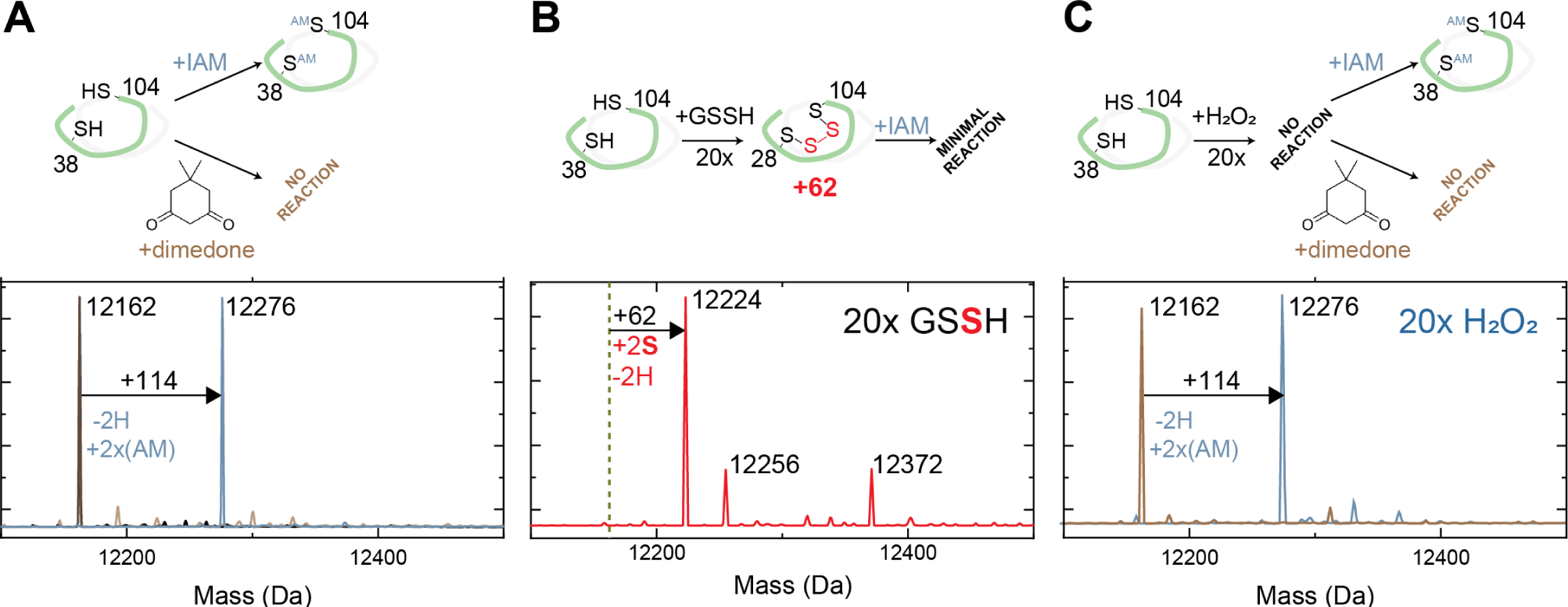
LC-ESI-MS analysis of HlyU *in vitro* reactivity upon the addition of GSSH or H_2_O_2_ **(A)** HlyU reacts with IAM but not with dimedone, showing that the protein is fully reduced. **(B)** HlyU reacts with GSSH to form a tetrasulfide link between its two cysteines, as seen in the mass shift of +62, corresponding to the addition of two sulfur atoms and the subtraction of two hydrogens. **(C)** HlyU does not react with H_2_O_2_, as seen by the absence of a peak corresponding to a dimedone adduct (115).

We further confirmed the specificity of persulfide-induced regulation by measuring the DNA binding affinities of *Vc*HlyU for a known DNA operator in the hemolysin promotor (11). These quantitative fluorescence anisotropy-based DNA binding experiments reveal that reduced HlyU binds to the operator upstream of HlyA with a 0.40 nM^-1^ affinity, which is comparable to other ArsR proteins. This affinity remains unchanged in the absence of reducing agent (Fig. 3A, Table 1). H_2_O_2_-pretreatment also leads to no change in DNA binding affinity, while GSSH-pretreatment yields a protein with virtually no DNA-binding activity (Fig. 3A-B). Moreover, upon titrating HlyU to saturation in absence of reducing agent it is possible to fully dissociate the protein from the DNA with the addition of a 10-fold excess of GSSH over HlyU thiol, as indicated by the decrease in anisotropy to that of the free DNA value. This dissociation is quite slow, however, consistent with the idea that reduction of the HlyU tetrasulfide may be protein-catalyzed in cells. DNA binding is only partially restored by addition of a reducing agent (Fig. 3C). In striking contrast, addition of 10-fold of H_2_O_2_ to DNA-bound HlyU leads to no change in anisotropy (Fig. 3D), providing further evidence that increased levels of H_2_O_2_ are not regulatory for DNA binding.

**Figure 3.**
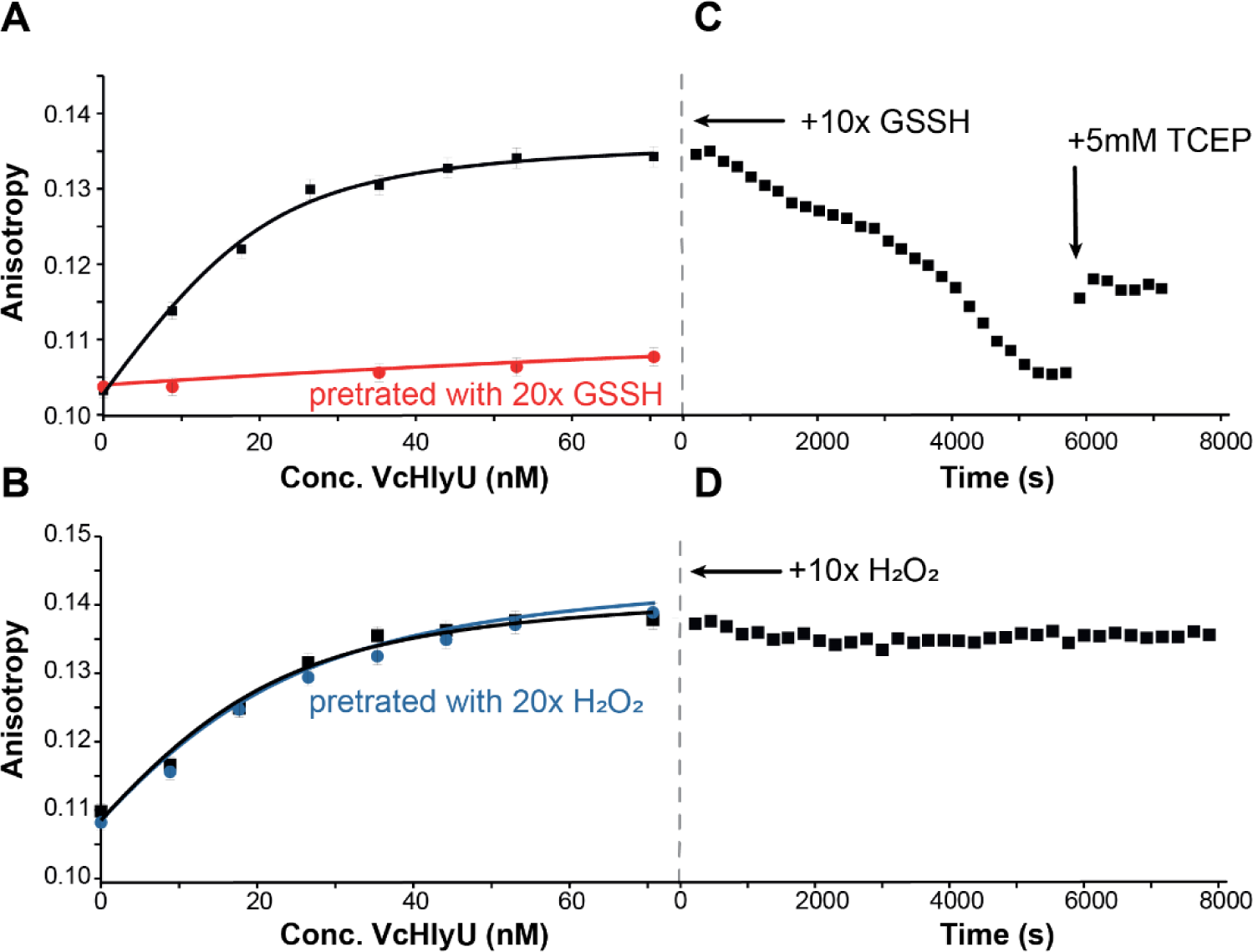
DNA binding isotherms of VcHlyU over its DNA operator in different oxidation states at 100mM NaCl: **(A)** reduced (black) vs. GSSH pre-treated (red) and **(A)** reduced (black) vs. H_2_O_2_ pre-treated (blue). Anisotropy changes of the fluorescein labeled HlyO operator with *Vc*HlyU after addition of a 10-fold excess of either (**C**) GSSH or (**D**) H_2_O_2_. After addition of oxidant anisotropy was followed over time until a new equilibrium condition was reached. Then a final concentration of 5mM TCEP was added to the solution to test reversibility of the oxidation.

**Table 1.** DNA binding affinities obtained for HlyU

| | $K_a$ [$\times 10^9 \text{ M}^{-1}$]<sup>a</sup> |
| --- | --- |
| **HlyU reduced** | $0.40 \pm 0.08$ ($n = 4$) |
| **HlyU, GSSH treated** | $0.009 \pm 0.005$ ($n = 2$) |
| **HlyU, H<sub>2</sub>O<sub>2</sub> treated** | $0.19 \pm 0.03$ ($n = 3$) |

<sup>a</sup>Errors correspond to standard deviations of the mean for n number of experiments carried out under the same conditions: 25 mM HEPES, pH = 7.0, 100 mM NaCl, 1 mM EDTA, 25°C. The treatment corresponds to a 5-fold addition relative to the protein subunit concentration.

Our MS-based reactivity assays and fluorescence anisotropy-based DNA binding experiments confirm that a post-translational thiol modification on HlyU negatively impacts DNA-binding and that these modifications are selective towards sulfur containing oxidants. To gain further insights on the structural impact of persulfide and peroxide treatment of HlyU homodimer, we used solution NMR as a probe for conformational changes that lead to DNA dissociation. ^1^H-^15^N HSQC spectra were acquired for HlyU dimer in the reduced, H_2_O_2_- and GSSH-treated states (Fig. S4). While the spectra obtained for HlyU in the reduced and H_2_O_2_-treated are essentially identical, GSSH treatment introduces significant perturbations in the spectrum, consistent with a well-folded dimer that is characterized by a distinct structure or distinct dynamics, which ultimately leads to DNA dissociation. To further explore the conformational changes induced by the tetrasulfide bond formation we performed a series of circular dichroism (CD) experiments at different temperatures to determine the stability of the secondary structure in the reduced, disulfide (diamide-treated (50)) and tetrasulfide crosslinked forms. The CD spectrum from HlyU is typical of a protein with significant secondary structure and a prevalence of α-helices, with a positive band <200 nm and negative signals at 210 and 218 nm (Fig. S5A). Reduced HlyU has a melting temperature (*T*_m_) of 65°C. Formation of the tetrasulfide decreases the native conformation stability by a small extent (*T*_m_ = 60 °C), while disulfide bond formation yields a far less stable form (*T*_m_ =52°C) (Fig. S5B). The comparatively lower stability of HlyU disulfide form is consistent with the observation of a high structural frustration in the disulfide-bonded structure of SqrR(50) and provides insights into the low reactivity of HlyU with oxidants that would ultimately lead to a disulfide-bonded form.

### Exogenous H_2_O_2_ and sulfide treatment of *Vibrio cholerae* cell cultures impair HlyU mediated p*hlyA* activation

Given that HlyU has properties of *bona fide* RSS-sensing repressor, we next aimed to determine the impact of cellular persulfides on HlyU-dependent regulation of hemolysin expression. Hemolysin expression is complex, and known to be regulated by HapR, Fur, and HlyU in *Vibrio cholerae* El Tor Serogroup O1 (11) (Fig. 4A). HlyU is somewhat unique among the ArsR family members in that instead of functioning as a repressor that inhibits the binding of RNA polymerase, it binds to the *hlyA* promoter and activates its transcription (65). The HlyU-dependent activation mechanism is best described in *Vibrio vulnificus* (24). Here, HlyU employs a “counter-silencing” mechanism, and displaces the global transcriptional repressor in Gram-negative bacteria, H-NS, from the two most upstream sites in the *hlyA* promoter, relieving transcriptional repression and leading to its activation upon HlyU recruitment. In *Vibrio cholerae,* the location of HapR, Fur, and HlyU binding has been described (65), and it has been shown that H-NS occupancy at the *hlyA* promoter is diminished by HlyU overexpression (82). However, the precise sites of H-NS binding are not known and the existing Chip-seq data suggest that H-NS may also bind coding regions (83). Thus, we first developed an assay where we could distinguish HlyU-mediated activation while decoupling hemolysin gene expression from quorum sensing and other environmental queues that impact these other transcriptional regulators involved in hemolysin regulation.

**Figure 4.**
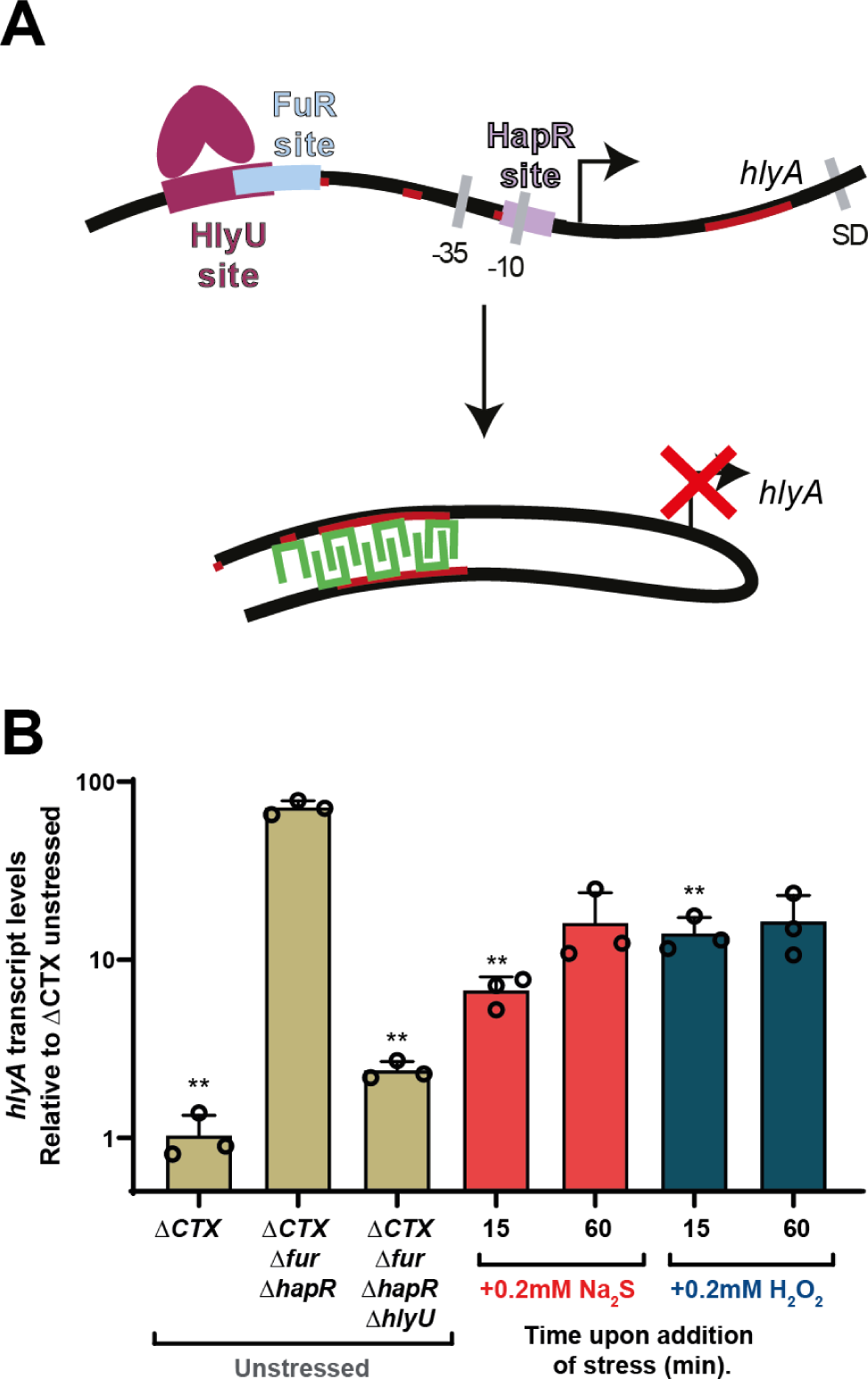
(**A**) Model of the mechanism of HlyU regulation of the *hlyA* gene, HlyU DNA dissociation leads to H-NS mediated repression. (**B**) Quantitative RT-PCR performed over a Δ*CTX* Δ*fur* Δ*hapR V. cholerae* strain with the addition of Na_2_S or H_2_O_2_. The bar chart shows the fold changes of induction of *Vc hlyA* after addition of Na_2_S (*red*) and H_2_O_2_ (*blue*), with transcript values normalized relative to the transcription level of *recA*. The values correspond to transcript levels relative to unstressed (UN) (middle *sand* bar) and are shown as mean ± SD from three replicate cultures. Statistical significance was established using a paired *t*-test relative to UN under the same conditions (**p<0.01, *p<0.05). Lines on the top of the chart show statistical significance relative to Δ*CTX* Δ*fur* Δ*hapR V. cholerae* mutant strain UN.

We first used a P_hlya_-*GFP* transcriptional reporter in different strain backgrounds to interrogate HlyU activity (Fig. S6). Consistent with previous findings (65), HlyU-mediated *hlyA* expression is highest in a *Δfur ΔhapR* background (Fig. S6A). This is also the case when *hlyA* transcript levels are followed by quantitative real-time PCR (qRT-PCR), and this activation is dependent on HlyU (Fig. 4A, *sand*-shaded bars). Furthermore, an H-NS deletion in the *Δfur ΔhapR* background provides additional support for the idea that the HlyU-dependent mechanism for P_hylA_ activation is eviction of H-NS (82) (Fig. S6B). However, it should be noted that the *Δhns* strains exhibit a very strong biofilm phenotype which complicates the fluorescence measurement (84). Interestingly, the difference in transcript levels of the *hlyA* and *gfp* genes that share the same P_hylA_ promotor suggest that HlyU-mediated activation may be enhanced by immediately adjacent regions, either downstream or upstream of the promotor, as the magnitude of transcriptional activation of *hlyA* is at least 20-fold higher in its native context relative to that of the *gfp* reporter, which only increases by 50% when HlyU is expressed (Fig. 4B and Fig. S7). This is not unexpected given the H-NS protection of P_hylA_ includes not only the promotor region but also at least 700 bp that encode for the HlyA protein (83).

We therefore elected to measure native *hlyA* transcript levels in a *Δfur ΔhapR* background, as it best recapitulates the H-NS eviction mechanism while isolating this event from Fur- and HapR-dependent effects. We next monitored HlyU-mediated transcriptional activation following acute Na_2_S and H_2_O_2_ stress (0.2 mM) in LB media at OD_600_≈0.2 after 15 and 60 min, as analogous persulfide sensors in other organisms are characterized by an acute phase transcriptional response (51). We used qRT-PCR to assess induction of *hlyA*, *gfp* (of the P_hlya_-GFP reporter) and *hlyU* mRNA expression. We observe a robust sulfide- and peroxide-inducible inhibition of the activation of *hlyA* expression (Fig. 4B, *red* and *blue* bars, respectively), while *gfp* and *hlyU* do not show significant differences upon stressor addition (Fig. S7). Exogenous treatment with these species is expected to trigger oxidative and sulfide stress signaling inside cells as they are both in equilibrium with membrane permeable species, H_2_S and neutral H_2_O_2_ respectively. Thus, our results suggest that an acute increase in intracellular levels of either H_2_S or H_2_O_2_ can downregulate *hlyA* transcription in a HlyU-mediated mechanism that likely involves H-NS polymerization on both coding and non-coding regions of the *hlyA* region. This crosstalk contrasts with the high degree of inducer specificity of HlyU suggested by the *in vitro* chemical reactivity, DNA-binding and NMR experiments. Nonetheless, our results show for the first time that HlyU-dependent activation of the *hlyA* operon can be modulated in response to exogenous stressors, including H_2_S.

### RSS metabolite profiling experiments

A change in the cellular levels of persulfidated low molecular weight thiols upon exogenous hydrogen sulfide stress is a robust biomarker for intracellular sulfide accumulation, having ultimately led to the identification of distinct features of thiols/RSS homeostasis in *S. aureus* (85), *E. faecalis* (37), *A. baumannii* (48), *Samonella* (86) and, more recently, *S. pneumoniae* (87) and *R. capsulatus*(88). Although the mechanistic details remain lacking, an emerging picture is that H_2_S is converted to thiol persulfides either via the enzymatic activity of SQR (89), or through other enzymatic (90) or non-enzymatic processes (91); this, in turn, leads to a transient increase in persulfidation of small molecule and proteome-derived thiols (90). The degree and identity of small molecule persulfidation, *i.e.,* speciation of the LMWT pool, depends to some extent on relative abundance of these species inside the cells (92). Changes in LMWT persulfidation have also been observed by endogenous production of H_2_S as deletion of the enzymes that biosynthesize it decreases the levels of persulfidation of LMWT, although the effect is small (48). The endogenous production of H_2_S can be also triggered by exogenous treatment with reactive oxygen species, such as H_2_O_2_. This H_2_O_2_-enhanced H_2_S endogenous production has been reported in *V. cholerae* and it has been shown that is has a critical role in cytoprotection against oxidative stress (47). However, the role of this H_2_S in signaling in *V. cholerae* is largely unknown, as is any quantitative information on LMWT thiols and RSS speciation in cells.

Given the paradoxical findings that HlyU does not react with H_2_O_2_ *in vitro*, yet H_2_O_2_ results in attenuation of HlyU-dependent activation, we hypothesized that both treatments trigger a change in reactive sulfur speciation that leads to HlyU modification. To test this hypothesis, we first employed an isotope dilution, electrophile trapping method to estimate the endogenous levels of the major cellular thiols, thiol persulfides, and inorganic sulfide and sulfane sulfur-bonded species in cell lysates from mid-exponential-phase cells (48) (Fig. 5A). Consistent with previous reports for *Vibrio* strains (93, 94) we find that cysteine and GSH are the major cellular thiols, followed by homocysteine at a concentration ≈5-10-fold lower (Fig. S8A). Basal H_2_S levels are much lower than that of the other thiols (Fig. 5B). Some HPE-IAM capped thiols and persulfuldes (*e.g.*, coenzyme A) are not easily quantified in the same chromatographic run and their concentrations were determined using monobromobimane (mBBr) as the capping agent (Fig. S9). We note that *V. cholerae* harbors complete glutathione (GSH) and coenzyme A (CoASH) biosynthetic pathways from cysteine (Fig. S8B), thus the level of intracellular thiols is expected to be comparable to other γ-proteobacteria (95, 96). The basal organic thiol persulfide-to-thiol ratio are all below 0.5%, as is the inorganic disulfide/sulfide ratio, findings generally consistent with other γ-proteobacteria and other bacteria that possess other LMWTs as the most abundant thiol (37, 48, 85, 86) (Fig. 5 and Fig. S9B).

**Figure 5.**
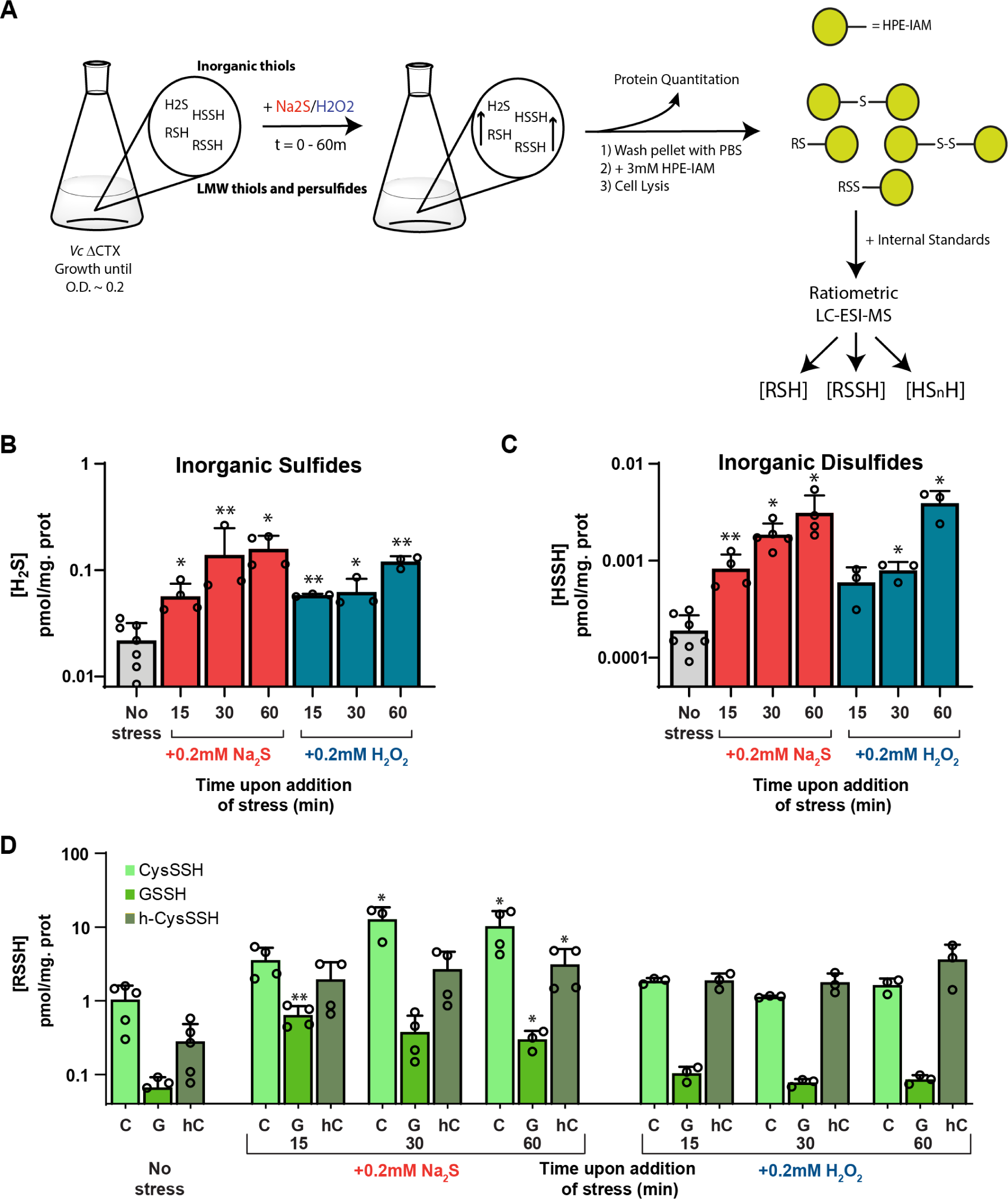
**(A)** Protocol scheme for LMW thiols and persulfide profiling. Growth of a ΔCTX *Vibrio cholerae* strain until O.D. of ∼0.2 is followed by the addition of Na_2_S or H_2_O_2_ to a final concentration of 0.2 mM. Cultures were centrifuged at 0 (prior to addition of stress), 15, 30 and 60 min. In all cases 1 mL of sample was withdrawn for protein quantitation. The metabolite profiling was generally carried out using HPE-IAM as labeling agent. The ratiometric LC-ESI-MS experiments were performed with the dilution of isotopically labelled internal standards of known concentration, which were used for quantitation of the organic and inorganic species. **(B)** Endogenous concentrations of hydrogen sulfide before and after addition of stress (Na_2_S, *red*; H_2_O_2_, *blue*) to mid-log-phase cultures. **(C)** Endogenous concentrations of hydrogen disulfide before and after addition of stress (Na_2_S, *red*; H_2_O_2_, *blue*) to mid-log-phase cultures. **(D)** Endogenous concentrations of cysteine (CysSSH), gluthathione (GSSH) and homocysteine (h-CysSSH) persulfides before and after addition of stress at different timepoints to mid-log-phase cultures. Statistical significance was established using a paired *t*-test relative to UN under the same conditions (**p<0.01, *p<0.05).

The addition of Na_2_S to *V. cholerae* cells leads to the expected transient increase on the GSSH levels (Fig. 5D and S10A) (37, 48), as well as a significant increase in the other organic and inorganic species detected in later time points (Figs. 5C and S10B-C). The addition of Na_2_S results also in a significant increase in cellular cysteine and homocysteine levels possibly as a result of increased flux through cysteine synthase (CysK) (Fig. S9B and S10A), with a corresponding increase in cysteine persulfide as well as H_2_S (Fig. 5B-C and S10C). Overall, our results suggest that the addition of exogenous sulfide results in its assimilation as organic thiol persulfides namely, GSSH, cysteine persulfide, as well as inorganic sulfane-sulfur bonded species, HSSH or S_2_. Strikingly, the addition of H_2_O_2_ results in an increase in intracellular sulfide and inorganic disulfide levels that nicely parallels that observed with exogenous Na_2_S treatment and is fully consistent with a previous study that showed that exogenous hydrogen peroxide treatment promotes endogenous H_2_S production (47)(Fig. 5B-C and S10). Interestingly, we observe little to no corresponding increase in the organic LMW thiol or thiol persulfide levels upon H_2_O_2_ treatment, at least for cysteine or glutathione (Figs. 5D and S10). This observation suggests that endogenous production of H_2_S does not necessarily have to impact the organic LMWT persulfide pool, a finding that suggests that HlyU is capable of sensing changes in either the inorganic or organic RSS pools. This result is consistent with prior findings with RSS sensors in other bacteria (50, 51); another possibility, not investigated here, is that HlyU reacts with protein persulfides or perhaps persulfidated H_2_S-producing enzymes themselves (90).

## DISCUSSION

In this work, we show that the *V. cholerae* hemolysin activator HlyU possess the characteristics of a *bona fid*e persulfide sensor and reacts exclusively with sulfane sulfur donors to form a tetrasulfide bridge between its two cysteines, which abrogates DNA-binding. While HlyU *in vitro* can distinguish between persulfides and other non-sulfur containing oxidants, exogenous treatment of *V. cholerae* cells with either sulfide (Na_2_S) or hydrogen peroxide (H_2_O_2_) prevents HlyU-dependent expression of the *hlyA* operon. By means of quantitative RSS metabolite profiling experiments, we show that sulfide (Na_2_S) or hydrogen peroxide (H_2_O_2_) increase the intracellular level of inorganic sulfane-sulfur bonded species, which likely triggers HlyU dissociation from the promoter and subsequent recruitment of H-NS to re-establish silencing of *hlyA* expression.

The remarkable functional diversity of the biochemically characterized ArsR family members in the persulfide sensor cluster (Cluster 4, Fig. 1) illustrates the widespread importance of (per)sulfide signaling in bacteria. On the one hand, in free living bacteria such as the purple bacterium *R. capsulatus*, persulfide sensing regulates sulfide-dependent photosynthesis (35, 88), while in plant symbionts and pathogens it is connected to biofilm formation and nodulation, shown in *Rhizobia* (71, 97). Human pathogens, such as *A. baumannii*, harness more than one persulfide-responsive regulator (the cluster 4 ArsR protein BigR and FisR, see Fig. 1A) and, in this way may connect sulfide metabolism with biofilm formation and metal homeostasis (48). One of the cluster 4 members in *E. coli* (YgaV) has been recently described as a master regulator connected to antibiotic resistance; however, its biochemistry remains unclear (98). In contrast, the enteric bacteria *Serratia marascens* encodes PigS, a cluster 4 candidate persulfide sensor that has been shown to regulate the production the red-pigmented antibiotic prodigiosin as well as sulfide metabolism genes (99). While PigS remains the sole biochemically characterized ArsR family protein with this function, antibiotic production regulation has also been shown for a known persulfide sensor from the CsoR family (100). These findings, that *bona fide* persulfide sensors regulate metabolic processes and pathways beyond sulfide detoxification, highlight the fact that sulfane sulfur species are not simply toxic molecules that bacteria need to metabolize and ultimately efflux, but function as reporters critical for survival in a particular microenvironment.

To what extent sulfide signaling is restricted to the three described families of dithiol-based transcriptional regulators is still a matter of debate, as the experimental strategies to interrogate the crosstalk between persulfides and other redox signaling molecules are still being developed. It is not yet clear whether a prototypical heme-based or cysteine-based redox sensor capable of responding to extracellular sulfide is indeed part of the global response towards these species (54–56). Most of the experiments performed here and in previous work suggest that even *bona fide* persulfide sensors such as SqrR (50, 88) and CstR (51) react slowly with LMW persulfides. These moderate *in vitro* rates contrast with the rapid transcriptional response that these species trigger in cells, raising the possibility that persulfide-specific regulators respond to persulfidated protein thiols instead of LMW persulfides (90). Addressing this question will provide new insights into the inducer specificity of thiol-based transcriptional regulators inside cells.

Moreover, it has been shown recently that bacterial (per)sulfide signaling is not necessarily restricted to thiol-based sensors since in *M. tuberculosis* (*Mtb*), the dormancy genes are regulated by a phosphorylation cascade in response to sulfide in a heme-dependent mechanism (101). Interestingly the expression of these genes is regulated by at least one of the ≈15 ArsR family proteins in *Mtb*. The considerable functional diversity of characterized ArsR proteins (SmtB (102), KmtR (103), CmtR (75), NmtR (104), Rv0081 (78), Rv2034 (72) and Rv2642(105)) coupled with sulfide-dependent regulation of dormancy genes and the possibility of more than one persulfide sensor suggests that bacteria such as *Mtb* may well incorporate a significant level of complexity or crosstalk between regulators. Thus, during infection, the signal triggered by different regulators that respond to persulfide, transition metal ion or peroxide signaling may be integrated into a single biological outcome (106, 107).

Pathogen colonization and disease progression are determined by major biochemical changes within the host during enteric infection (33). Successful pathogens must sense and respond to these biochemical changes, that may be also accompanied by dramatic changes in the microbiota, particularly induced by antibiotics (106). In the case of *Vibrio* spp. the response to the biochemical changes in the gut plays a crucial role in exotoxin expression (1, 47). Here, we show that changes in intracellular persulfides in response to changes in oxidative stress and sulfide stress prevalent in the inflamed gut (32, 33, 47) can attenuate hemolysin expression. This link between persulfide sensing and other adaptive responses at the host-pathogen interface may help understand how other pathogens or pathobionts respond to changes in the gut or other infected tissues where these species are prevalent and/or biosynthesized. Given the importance of (per)sulfide signaling, we expect that more of these *bona fide* persulfide sensing proteins will continue to be discovered or rediscovered orchestrating distinct adaptative responses in bacteria.

## EXPERIMENTAL PROCEDURES

### *V. cholerae* strains, growth media

All *V. cholerae* strains used in this study are derivatives of strain E7946 (107), see Table S1. Mutant constructs were generated via splicing by overlap extension PCR (SOE PCR) and introduced into cells by chitin-dependent natural transformation exactly as previously described (108). Routine culture employed Luria-Bertani medium at 30°C for overnight growth, and 37°C with constant agitation. When appropriate, culture medium was supplemented with trimethoprim (10 µg/mL), carbenicillin (20 µg/mL), kanamycin (50 µg/mL), spectinomycin (200 µg/mL), and/or chloramphenicol (1µg/mL).

### Protein preparation

*Vc*HlyU was subcloned into a pHis plasmid with NcoI and NedI encoding the untagged protein. *Vc*HlyU was expressed in *E. coli* BL21(DE3) cells at 16°C overnight after induction with 1 mM IPTG. Freshly collected cells expressing *Vc*HlyU were suspended in 120 mL of buffer B (25 mM MES, 750 mM NaCl, 2 mM tris(2-carboxyethyl)phosphine (TCEP), 1 mM EDTA, pH 6.0) and lysed by sonication using a Fisher Scientific model 550 sonic dismembrator. The cellular lysate was centrifuged at 8,000 r.p.m. for 15 min at 4 °C. The supernatant was collected and subjected to protein and nucleic acid precipitation by addition of 10% polyethylenimine (to 0.015% v/v) at pH 6.0. After stirring for 1 h at 4 °C, the solution was clarified by centrifugation at 8,000 r.p.m. for 15 min at 4 °C, with the supernatant precipitated by the addition of (NH_4_)_2_SO_4_ to 70% saturation with stirring for 2 h. After centrifugation at 8,000 rpm for 15 min, the precipitated protein was dissolved and dialyzed against buffer A (25 mM MES, 150 mM NaCl, 2 mM TCEP, 1 mM EDTA, pH 6.0). This solution was loaded onto a 10-mL SP (sulfopropyl) Fast Flow cation exchange column equilibrated with buffer A. The protein was then eluted using a 150-mL linear gradient from 0.150 to 0.75 M NaCl. *Vc*HlyU samples for NMR experiments were isotopically labeled using isotopes for NMR experiments purchased from Cambridge Isotope Laboratories. Cells were grown in an M9 minimal media containing (per liter of growth media): 6g Na_2_HPO_4_, 3g KH_2_PO_4_, 0.5g NaCl, 0.24g MgSO_4_, 0.011g CaCl_2_, 1mg of thiamine, 2g of ^13^C-glucose, 0.5g ^15^N-NH_4_Cl and 1/1000 ampicillin until an O.D. of 0.6 was reached. Induction, expression, and purification conditions were the same as the above protocol. All the proteins characterized here eluted as homodimers, as determined by a calibrated Superdex 75 (GE Healthcare) gel filtration chromatography column (25 mM MES, 0.2 M NaCl, 2 mM EDTA, 2 mM TCEP, 5% glycerol, pH 6.0, 25 °C).

### Fluorescence anisotropy-based DNA binding experiments

Standard fluorescence anisotropy-based DNA binding experiments were carried out using two 29-base-pair fluorescein (F)-labeled operator DNA fragments, 5’F-(T) AT AAA TTA ATT CAG ACT AAA TTA GTT CAA A-3’ and its complement: 5’-TTT GAA CTA ATT TAG TCT GAA TTA ATT TAT A-3’ from the hlyA promotor region (HlyO), in DNA binding buffer (10 mM HEPES, pH 7.0, 0.1 M NaCl, in presence or in absence of 2 mM TCEP). After the final addition of protein, *Vc*HlyU was either dissociated from the DNA by addition of an excess of GSSH or H_2_O_2_ (when performed in the absence of reducing agent) or associated with the DNA with the addition of sufficient (5 mM) TCEP to reduce the tetrasulfide-crosslinked HlyU. This experiment validates that the tetrasulfide formation is partially reversible and inhibitory to the association of HlyU with the DNA. All experiments were performed in triplicate. All anisotropy-based data were fit to a simple 1:1, nondissociable dimer binding model to estimate *K*_a_ using DynaFit(109).

### LC-ESI-MS analysis of derivatized proteins

Reduced *Vc*HlyU was buffer exchanged anaerobically into a degassed 300 mM sodium phosphate buffer, pH 7.4, containing 1 mM EDTA. Reactions containing 60 µM protein were anaerobically incubated at room temperature with a 20-fold excess of oxidizing reagent; namely GSSH or H_2_O_2_ for 1 h or for different times as indicated in the figures. The reactions were quenched by addition of equal volumes of 60 mM IAM in the case of the RSSH or 60 mM of 5,5-dimethylcyclohexane-1,3-dione (dimedone) in the case of H_2_O_2_. Analysis was performed in the Laboratory for Biological Mass Spectrometry at Indiana University using a Waters Synapt G2S mass spectrometer coupled with a Waters ACQUITY UPLC iClass system. Protein samples of 5 µL were loaded onto a self-packed C4 reversed-phased column and chromatographed using an acetonitrile-based gradient (solvent A: 0% acetonitrile, 0.1% formic acid; solvent B: 100% acetonitrile, 0.1% formic acid). Data were collected and analyzed using MassLynx software (Waters).

For aerobic conditions, *Vc*HlyU was exchanged into a 25 mM MES pH 6.0, 150 mM NaCl containing 1 mM EDTA. Reactions containing 60 µM protein were aerobically incubated at room temperature with a 20-fold excess of oxidizing reagent; namely GSSH or H_2_O_2_ for 1 h or overnight, as indicated in the figures. The reactions were quenched by addition of equal volumes of 60 mM IAM or dimedone. Protein preparations were sent to the Proteomics Core Facility of CEQUIBIEM at the University of Buenos Aires. Samples were desalted with Zip-Tip C18 (Merck Millipore), and proteins were analyzed with Orbitrap technology (Q-Exactive with High Collision Dissociation cell and Orbitrap analyzer, Thermo Scientific, Germany). Peptide Ionization was performed by ESI. Data was analyzed with the Xcalibur Software from ThermoScientific.

### Preparation of RSSH

GSSH, CysSSH and h-CysSSH were freshly prepared by mixing a five-fold molar excess of freshly dissolved Na_2_S with RSSR and incubating anaerobically at 30 °C for 30 min in degassed 300 mM sodium phosphate (pH 7.4). The concentration of the *in situ*-generated persulfides were determined using a cold cyanolysis assay as previously described(37), and was used without further purification at the indicated final concentrations (88).

### P_hylA_-GFP fluorescence measurements

Strains were grown overnight in plain LB medium. Cells were then washed and concentrated in instant ocean medium (7 g/L; Aquarium Systems) to an OD_600_ = 5.0. Then, 200 µ L for each sample was transferred to a 96-well plate and GFP fluorescence was measured on a Biotek H1M plate reader. The background fluorescence was determined by using a wildtype E7946 strain (that lacks GFP), and subtracted from all samples.

### Quantitative real time PCR analysis

*V. cholerae* strains (Table S1) were inoculated from glycerol stocks into 5 mL LB medium and grown at 30 °C overnight. The overnight culture was diluted 1/100 into 15 mL LB medium at a starting OD600 ≈ 0.002, grown to an OD600 of 0.2 at 37 °C followed by the addition of stressor, Na_2_S (0.2 mM), or H_2_O_2_ (0.2 mM). An aliquot of the cultures was centrifuged for 10 min prior to the addition of stressor (*t*=0), as well as 15 min and 60 min post addition of stressor to the cultures. Following centrifugation, the cell pellets were washed twice with ice-cold PBS, centrifuged for 5 min and stored at –80 °C until use. Pellets were thawed on ice and resuspended in 1 mL of TRI Reagent (catalog no. TR-118; Molecular Research Center). Resuspended cells were placed in tubes containing 0.1-mm silica beads (Lysing matrix B tubes, catalog no. 6911-100; MP Biomedicals) and lysed in a bead beater (Bead Ruptor 24 Elite; Omni) at a rate of 6 m/s for 45 s twice, with a 5-min cooling on ice between runs. Then, 200 µL of chloroform was added, followed by vigorous mixing and centrifugation for 15 min at 13,200 rpm. The top aqueous layer was removed to a new tube, and 70% ethanol was added at a 1:1 volume ratio. RNA purification was completed using the RNeasy minikit (catalog no. 74104; Qiagen) following DNase I treatment (catalog no. 79254; Qiagen). Next, 5 µg of total RNA was subsequently digested with the DNA-free kit (catalog no. AM1906; Ambion) and diluted 5-fold. First-strand cDNA was synthesized using random hexamers (Quanta Biosciences) and a qScript Flex cDNA synthesis kit (catalog no. 95049-100; Quanta Biosciences). Reactions contained 10 µL of 2× Brilliant III Ultra-Fast SYBR green QPCR master mix (catalog no. 600882; Agilent), 2 µL each of 2 µM PCR primers (see Table S1 for used primers), 0.3 µL of 2 µM ROX reference dye, and 6 µL of diluted cDNA. Relative transcript amounts were measured using the MX3000P thermocycler (Stratagene) running the SYBR Green with dissociation curve program and normalized to the amount of *recA*. The thermal profile contained 1 cycle at 95 °C for 3 min, 40 cycles at 95 °C for 20 s to 59 °C for 20 s. Subsequently, a dissociation curve starting at 55 °C going to 95 °C in 0.5 °C increments with a dwell time of 30 s was performed to assess the specificity of the reactions. Three biologically independent samples were measured for each treatment and the mean ± standard deviation (S.D.) values are reported.

### Quantitation of cellular LMW thiols and persulfides

Overnight ΔCTX *V. cholerae* cells grown in LB media were diluted to an OD600 of 0.02 in LB and grown in sealed tubes with constant agitation at 37 °C. When these cultures reached an OD600 of 0.2, 0.2 mM disodium sulfide or H_2_O_2_ was added. Samples were collected before (*t*=0 min) addition of the stressor and at 15, 30 and 60 min following addition of stressors. They were centrifuged at 3,000 rpm for 10 min. The resulting pellets were washed with ice-cold PBS, pelleted again by centrifugation (16,100 rpm for 5 min), and stored frozen at −80°C until use. Thawed cell pellets were resuspended in 200 µL of a PBS solution containing 3mM of b-hydroxyphenyl iodoacetamide (HPE-IAM) labeling agent, or in 100 µL of mBBr solution containing 20 mM TRIS-HBr, pH 8.0, 50% acetonitrile, 1 mM mBBr. The resuspension solutions were then subjected to three freeze-thaw cycles in liquid nitrogen in the dark. Cell debris was removed by centrifugation, the supernatant was transferred to a 0.2 µm-pore-size centrifugal filter unit and centrifuged at 13200 for 10 min prior to injection into a liquid chromatograph mass spectrometry (LC-MS) system for quantitation of LMW thiol and persulfides as follows.

Samples (10 µL) were injected into a Triart C18 column (YMC, Inc.) (50 by 2.0 mm inner diameter) and subjected to chromatography on a Waters Acquity Ultra Performance Liquid Chromatography (UPLC) I-class system, using a methanol-based gradient system (for solvent A, 10% methanol and 0.25% acetic acid, pH 3.0; for solvent B, 90% methanol and 0.25% acetic acid, pH 3.0) with the elution protocol at 25°C and a flow rate of 0.2 ml/min as follows: at 0 to 3 min, 0% B isocratic; at 3 to 7 min, 0% to 25% B, linear gradient; at 7 to 9 min, 25% B isocratic; at 9 to 12 min, 25% to 75% B, linear gradient; at 12 to 14 min, 75% to 100% B, linear gradient; at 14 to 14.5 min, 100% B isocratic, followed by re-equilibration to 0% B. Quantitation of LMW thiols and persulfides was carried out with a Waters Synapt G2S mass spectrometer by spiking in a specific amount of authentic LMW persulfide standards (HPE-IAM heavy) synthesized with deuterium isotopic labelling. The persulfide standards used for quantification were GSSH, CysSSH and h-CysSSH for their respective thiols and persulfides, and h-CysSSH for quantification of CoASH, CoASSH (mBBr labeling) and of inorganic sulfides and disulfides. Analysis of peak areas was performed in Masslynx (v 4.1) software, and the data were normalized to protein concentrations measured using a Bradford assay with bovine serum albumin (BSA) as the standard, as previously described(38). Data shown represent means and standard deviations of results from at least three biological replicates.

### Sequence Similarity Network (SSN) Analysis

The EFI-EST webserver (https://efi.igb.illinois.edu/efi-est/) was used to generate a sequence similarity network (SSN) using the Pfam PF01022 and Interpro IPR001845 databases of ArsR proteins as input, using the “Families” option of the webserver(74). The initial computation parameters were left at their default values, with an E-Value of 5. Given the large number of sequences used as input (365,837), an UniRef50 database was generated, where each node in the SSN groups sequences that share at least 50% of sequence identity. For the final calculation of the network, sequences shorter than 50 residues in length or longer than 150 residues were left out of the network, as such short sequences likely come from peptide fragments and those longer than 150 residues cannot correspond to the canonical ArsRs, which have only one domain(70). For the final network, an Alignment Score of 22 was used as threshold to cluster together UniRef50 nodes with at least 40% of sequence identity. The SSN was visualized using the Cytoscape software, version 3.8.0(110). The network was represented using an unweighted prefuse force-directed layout. The network was colored, and the sequence logos and further cluster analysis were done using the “SSN utilities” of the EFI-EST webserver (74). Previously biochemically characterized ArsR (Table S2) were found in the network by manually searching for their Uniprot ID in Cytoscape. A Genome Neighborhood Analysis was performed in the EFI-GNT webserver(74) using sub-networks of the main network (Fig S1B) form by each of the main (sub)clusters as input, with a neighborhood size of 5 and a minimal co-occurrence percentage lower limit of 5. Sequence logos for each cluster were obtained by performing a multiple sequence alignment (MSA) of the UNIREF50 sequences from each (sub)cluster using the MEGA software (version 11)(111) and the MUSCLE algorithm(112). The gap extension penalty was set to −0.5 and the neighbor-joining algorithm was selected as a cluster method in all iterations. All other parameters for the MSA were left in their default values. Alignments were manually curated by removing portions of the N-terminal and C-terminal in abnormally long sequences. Gaps accounting for indels in a minority of the sequences aligned were removed to avoid the overestimation of the conservation of residues in those positions in the sequence logos. Logos were generated using the Skylign web server(113) with default parameters using the MSAs exported from MEGA as inputs. The secondary structure prediction for the sequence logos shown in Fig. 1B was obtained using the JPred4 webserver(114), using the consensus sequence for each cluster obtained in their MSA. All the consensus sequences from the studied clusters shared the same secondary structure prediction. Sequences within certain clusters in the network were assigned a putative function based on the information provided by their genome neighborhood, the conservation of key sequence motives within a cluster such as the DNA-binding residues(66) or the ligand-binding residues(70), and the regulatory function of characterized ArsRs that mapped within that cluster.

### NMR spectroscopy

Typical NMR sample solution conditions were 200 µM ^15^N-labeled wild-type HlyU in 20 mM MES pH 6, 250 mM NaCl, 1 mM EDTA buffer as indicated. A Bruker 600 MHz spectrometer equipped with room temperature probe was used to acquire data for all HlyU samples. NMR data were processed using NMRPipe and were analyzed using Sparky. All spectra were acquired at 30 °C as indicated. Chemical shift is referenced relative to 2,2-dimethyl-2-silapentene-5-sulfonic acid (DSS).

### CD spectroscopy

The CD spectra were recorded at 25 °C using a JASCO-810 spectropolarimeter flushed with N_2_ and a 0.1 cm path length cuvette. Ten spectra were registered from 300 to 195 nm, at 0.1 nm intervals. Unfolding of secondary structure was followed by heating a protein solution (33 µM) from 25 to 90 °C and following the decrease in ellipticity at 235 nm using a 1.00 cm path length cuvette. A 25 mM HEPES, 200 mM NaCl and 1 mM TCEP (reducing agent only added to the solution for the experiment in reduced HlyU) buffer was used in these experiments.

## DATA AVAILABILITY

The authors declare that all data supporting the findings of this study are available within the paper and its Supplementary information files. All raw data are available from the corresponding author upon request.

## ACKNOWLEDGEMENTS

We gratefully acknowledge support by the MinCyT Argentina (PICT 2019-0011, 2019-3805 to D.A.C.), US National Institutes of Health (R35 GM118157 to D.P.G. and R35 GM128674 to A.B.D.). D.A.C. is a Staff Member of CONICET, Argentina; C.P.D. and G.T.A. are supported by a postdoctoral and doctoral fellowship provided by CONICET, Argentina, respectively; and G.T.A was also supported by the PROLAB program of the American Society of Biochemistry and Molecular Biology (ASBMB).

