## Supporting Information for "Increased intracellular persulfide levels attenuate HlyU-mediated hemolysin transcriptional activation in *Vibrio cholerae*"

**This file contains Supporting Tables S1-S2, Supporting Figures S1-S10.**

**Table S1:** *V. cholerae* strains and primers used for qRT PCR used for this study.

| Strain | Description |
|---|---|
| TND0004 | *WT* |
| TND2438 | $\Delta lacZ::P_{hlyA}$-*msfGFP* $Cm^R$ |
| TND2455 | $\Delta hlyU::Kan^R$, $\Delta lacZ::P_{hlyA}$-*msfGFP* $Cm^R$ |
| TND2459 | $\Delta hapR::Spec^R$, $\Delta lacZ::P_{hlyA}$-*msfGFP* $Cm^R$ |
| TND2456 | $\Delta hlyU::Kan^R$, $\Delta hapR::Spec^R$, $\Delta lacZ::P_{hlyA}$-*msfGFP* $Cm^R$ |
| TND2533 | $\Delta fur::Tm^R$, $\Delta hapR:Spec^R$, $\Delta lacZ::P_{hlyA}$-*msfGFP* $Cm^R$ |
| TND2534 | $\Delta fur::Tm^R$, $\Delta hapR::Spec^R$, $\Delta hlyU::Kan^R$, $\Delta lacZ::P_{hlyA}$-*msfGFP* $Cm^R$ |
| TND2531 | $\Delta fur:Tm^R$, $\Delta lacZ::P_{hlyA}$-*msfGFP* $Cm^R$ |
| TND2532 | $\Delta fur:Tm^R$, $\Delta hlyU:Kan^R$, $\Delta lacZ::P_{hlyA}$-*msfGFP* $Cm^R$ |
| TND3182 | $\Delta hapR:Spec^R$, $\Delta fur::Tm^R$, $\Delta lacZ::P_{hlyA}$-*msfGFP* $Cm^R$ |
| TND3183 | $\Delta hapR:Spec^R$, $\Delta fur::Tm^R$, $\Delta hns::Carb^R$, $\Delta lacZ::P_{hlyA}$-*msfGFP* $Cm^R$ |
| TND3185 | $\Delta hapR:Spec^R$, $\Delta fur::Tm^R$, $\Delta hlyU::Kan^R$, $\Delta lacZ::P_{hlyA}$-*msfGFP* $Cm^R$ |
| TND3186 | $\Delta hapR:Spec^R$, $\Delta fur::Tm^R$, $\Delta hlyU::Kan^R$, $\Delta hns::Carb^R$, $\Delta lacZ::P_{hlyA}$-*msfGFP* $Cm^R$ |
| **Primers PCR** | **Sequence** |
| *recA* – rev | GCGCAGCAATCTTGTTCTTC |
| *recA* – fwd | CGTTTGGATATTCGCCGTACT |
| *hlyA* – rev | CTC TGT GGC TGA GGC TTT AT |
| *hlyA* – fwd | CGA TGC TTT GTG GGT GAA TAC |
| *GFP* – rev | GCTCTTGCACGTATCCTTCT |
| *GFP* – fwd | TTGTGACGACTCTGACTTATGG |
| *hlyU* – rev | CCA CGC TAG ATG TTG AGA AAG A |
| *hlyU* – fwd | GGA CAA TGA ACT GTC GGT AGG |

**Table S2:** Characterized ArsRs in the SSN.

| Protein Name* | Organism | Uniprot ID | Cluster number in Network |
|---|---|---|---|
| ecArsR (116) | *Escherichia coli* | P37309 | 1A |
| ArsR2 (117) | *Escherichia coli* | A0A142BMN6 | 1A |
| ArsR(118) | *Staphylococcus aureus* | P30338 | 1A |
| ArsR1&2 (119) | *Geobacillus kaustophilus* | Q5KUX7 | 1A |
| ArsR (120) | *Pseudomonas putida* | Q88LK1 | 1A |
| AseR(67) | *Bacilus subtilis* | P96677 | 1A |
| ArsR (77) | *Corynebacterium glutamicum* | A0A5H1ZR36 | 1A |
| Rv2642(105) | *Mycobacterium tuberculosis* | P71941 | 1A |
| AztR(121) | *Cyanobacterium anabaena* | Q8ZS91 | 1B |
| BxmR(122) | Oscillatoria brevis | Q76L30 | 1B |
| NmtR(104) | *Mycobacterium tuberculosis* | O69711 | 1B |
| ZiaR(123) | *Synechocystis sp.* | Q55940 | 1B |
| CadC(124) | *Staphylococcus aureus* | P20047 | 1B |
| SmtB(125) | *Synechococcus elongatus* | P30340 | 1B |
| CzrA(126) | *Staphyloccocus aureus* | O85142 | 1B |
| CzrA(67) | *Bacilus subtilis* | O31844 | 1B |
| CadC(127) | *Listeria innocua serovar 6* | P0A4U2 | 1B |

| | | | |
|---|---|---|---|
| SmtB(102) | *Thermus thermophilus* | Q72KG0 | 1B |
| CadC (127) | *Lysteria monocytogenes* | Q56405 | 1B |
| Rv2034(72) | *Mycobacterium tuberculosis* | O53478 | 2 |
| SdpR (128) | *Bacilus subtilis* | O32242 | 2 |
| Rv0081(78) | *Mycobacterium tuberculosis* | P9WMI7 | 3 |
| AntR(129) | *Comamonas testosteroni* | A0A096FLR2 | 3 |
| BigR(48) | *Acinetobacter baumanni* | D0C7U0 | 4 |
| YgaV(130) | *Escherichia coli* | P77295 | 4 |
| NolR(71) | *Rhizobium fredii* | Q83TD2 | 4 |
| SqrR(50) | *Rhodobacter capsulatus* | D5AT91 | 4 |
| HlyU(11)** | *Vibrio cholerae serotype O1* | P52695 | 4 |
| HlyU(20) | *Vibrio parahaemolyticus* | Q87S95 | 4 |
| HlyU(19) | *Vibrio vulnificus* | A0A3Q0L222 | 4 |
| BigR(40) | *Xylella fastidiosa* | Q9PFB1 | 4 |
| BigR(131) | *Agrobacterium tumefaciens* | Q8UAA8 | 4 |
| PigS (99) | *Serratia* sp. strain ATCC 39006 | E7BBJ0 | 4 |
| SoxR(132) | *Pseudaminobacter salicylatoxidans* | Q5ZQN5 | 4 |
| ArsR(77) | *Acidithiobacillus ferrooxidans* | B7J952 | 5 |
| ArsR(133) | *Agrobacterium tumefaciens ArsR 1* | H0HHH0 | 5 |
| CyeR(134) | *Corynebacterium glutamicum* | A4QI86 | 6 |
| YczG(135) | *Bacilus subtilis* | O31480 | 6 |
| RexT(61) | Nostoc sp. | Q8YVV6 | 6 |
| MerR(136) | *Streptomyces lividans* | P30346 | 8 |
| PyeR(137) | *Pseudomonas aeruginosa* | Q9HW47 | 9 |
| KmtR(103) | *Mycobacterium tuberculosis* | O53838 | 10 |
| PagR(138) | *Bacillus anthracis* | O31178 | 14 |
| SmtB(139) | *Mycobacterium tuberculosis* | P9WMI4 | 16 |
| SrnR(73) | *Streptomyces griseus* | Q8L1Y3 | 21 |
| CmtR(75) | *Streptomyces coelicolor* | Q9RD34 | 22 |
| CmtR(140) | *Mycobacterium tuberculosis* | P9WMI8 | 22 |

* The protein name is followed by the most updated publication with the available biochemistry information. ** This work.

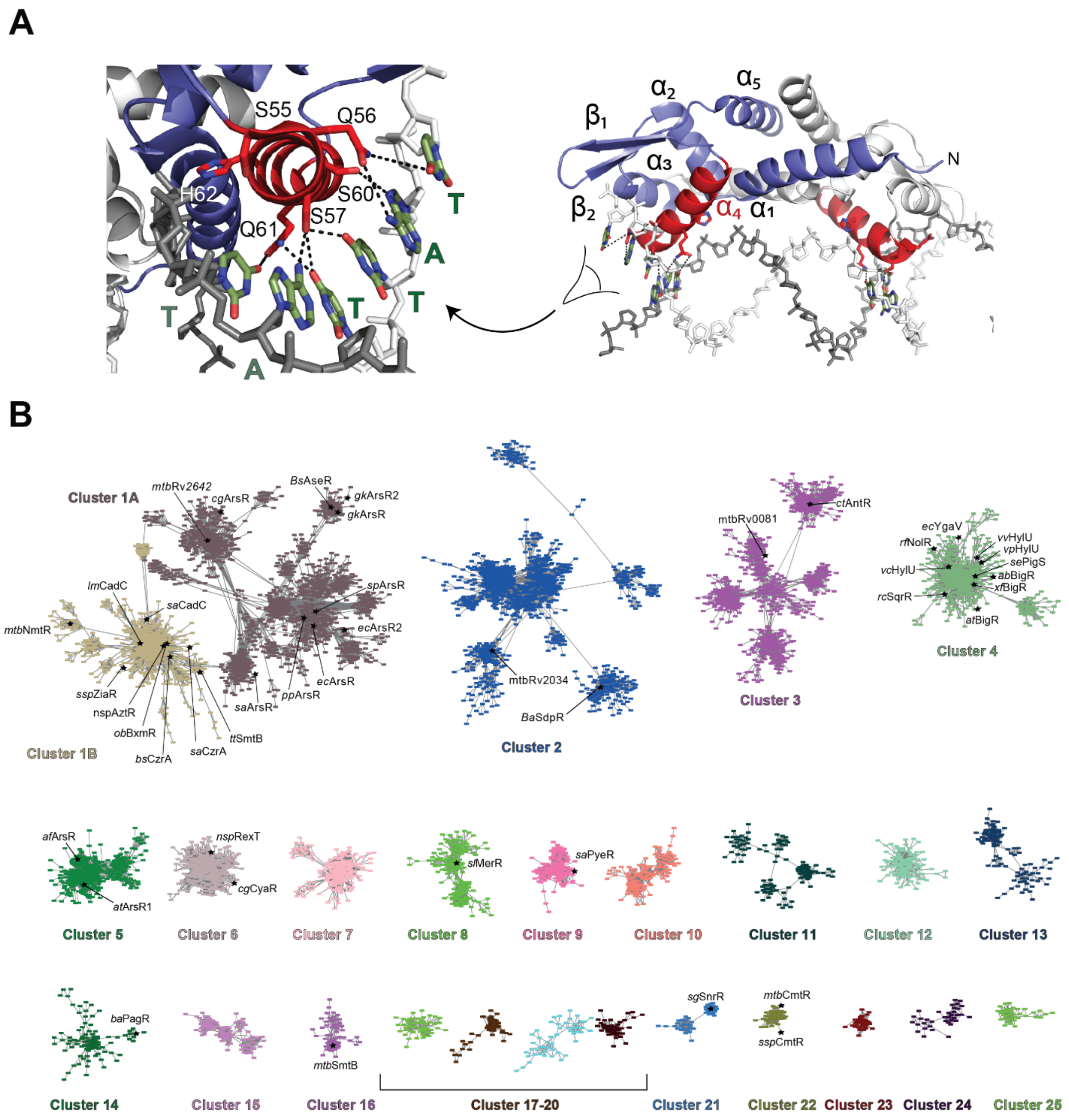


**Figure S1. (A)** ArsR superfamily proteins architecture (a1-a2-a3-a4-b1-b2-a5) and DNA recognition exemplified by the structure of DNA-bound NolR. The contacts between the protein sequence in a4 (*red*) and bases from DNA (*green*) in the first half-site are marked with *black* dashes. **(B)** All the main clusters generated in the sequence similarity network obtained using Pfam PF01022 and Interpro IPR001845 datasets of annotated ArsRs proteins.

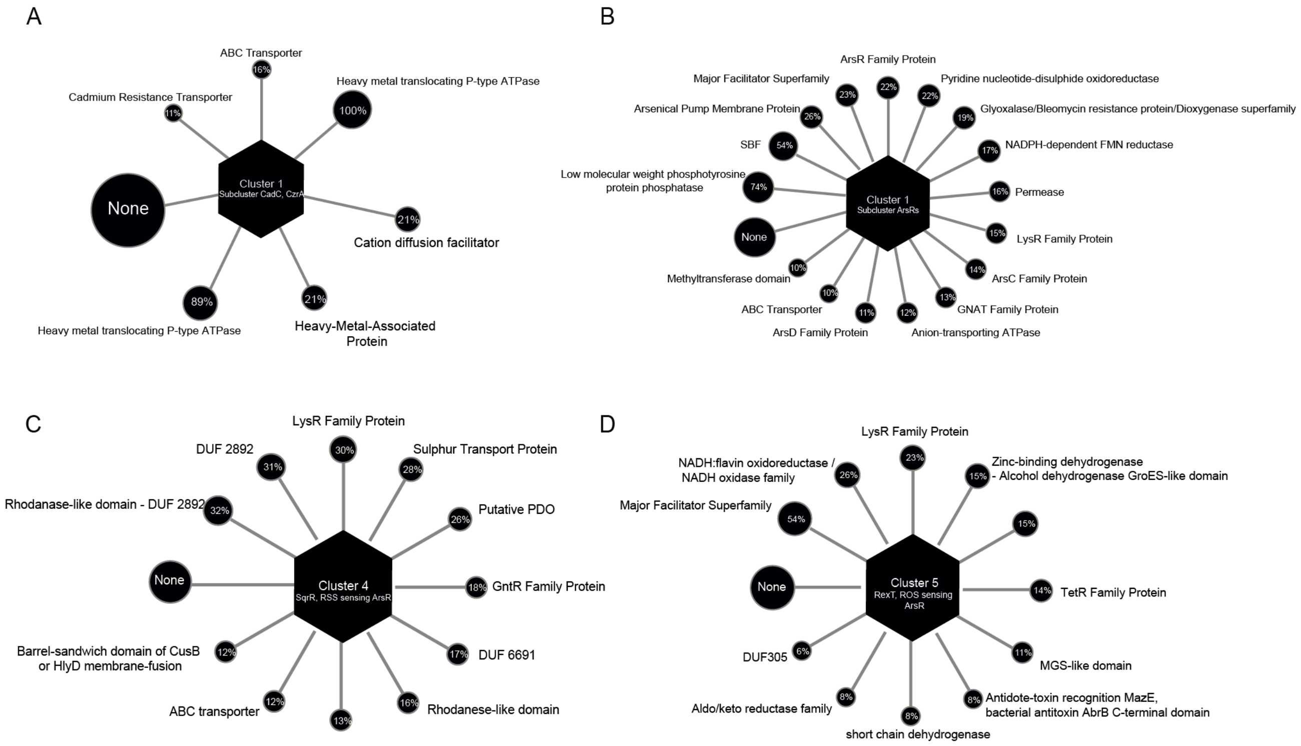


**Figure S2.** Genomic neighborhood and location of ArsR genes from cluster 1 (A and B), cluster 4 (C) and cluster 5 (D).

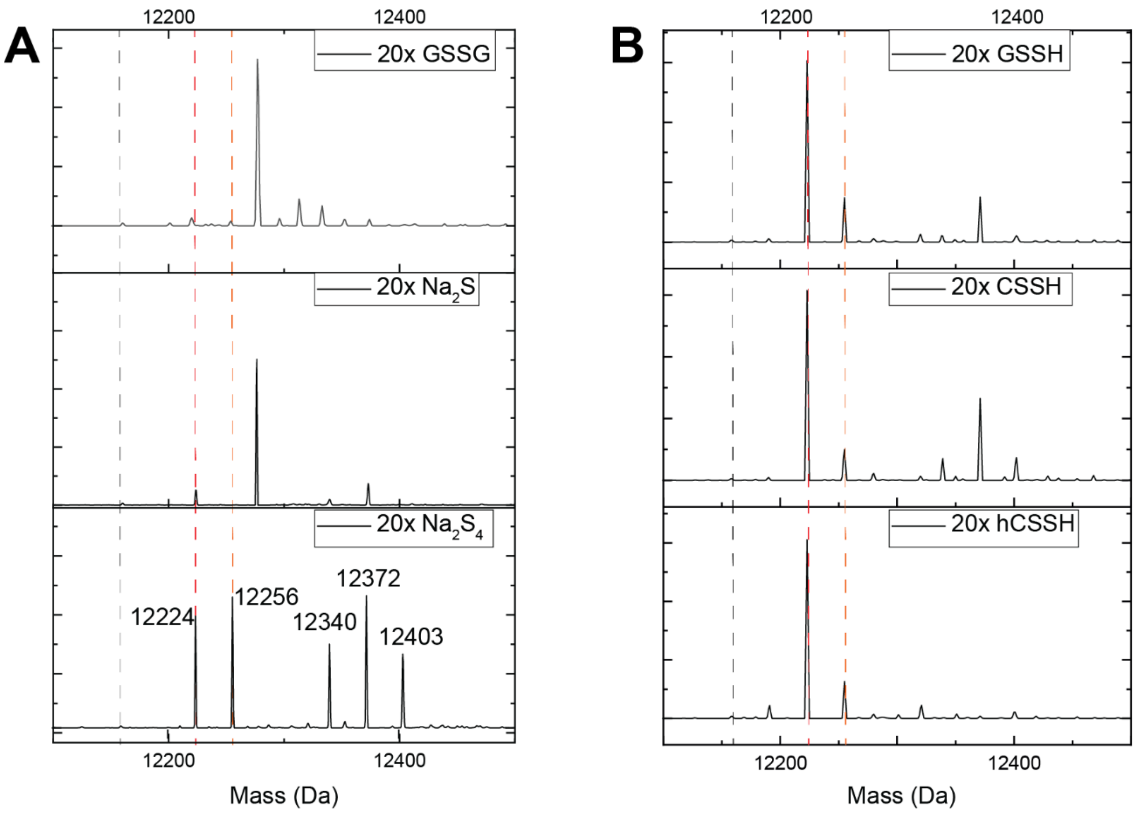


**Figure S3.** LC-ESI-MS analysis of HlyU *in vitro* reactivity upon a one-hour incubation with a 20-fold excess of **(A)** GSSG, $Na_2S$ and $Na_2S_4$ and **(B)** organic persulfides GSSH, cysteine and homocysteine persulfides (CSSH and hCSSH, respectively) and then capped with IAM. *Grey* dashed lines correspond to the reduced and uncapped HlyU monomer, while the two *red* dashed lines correspond to the tetrasulfide and pentasulfide species.

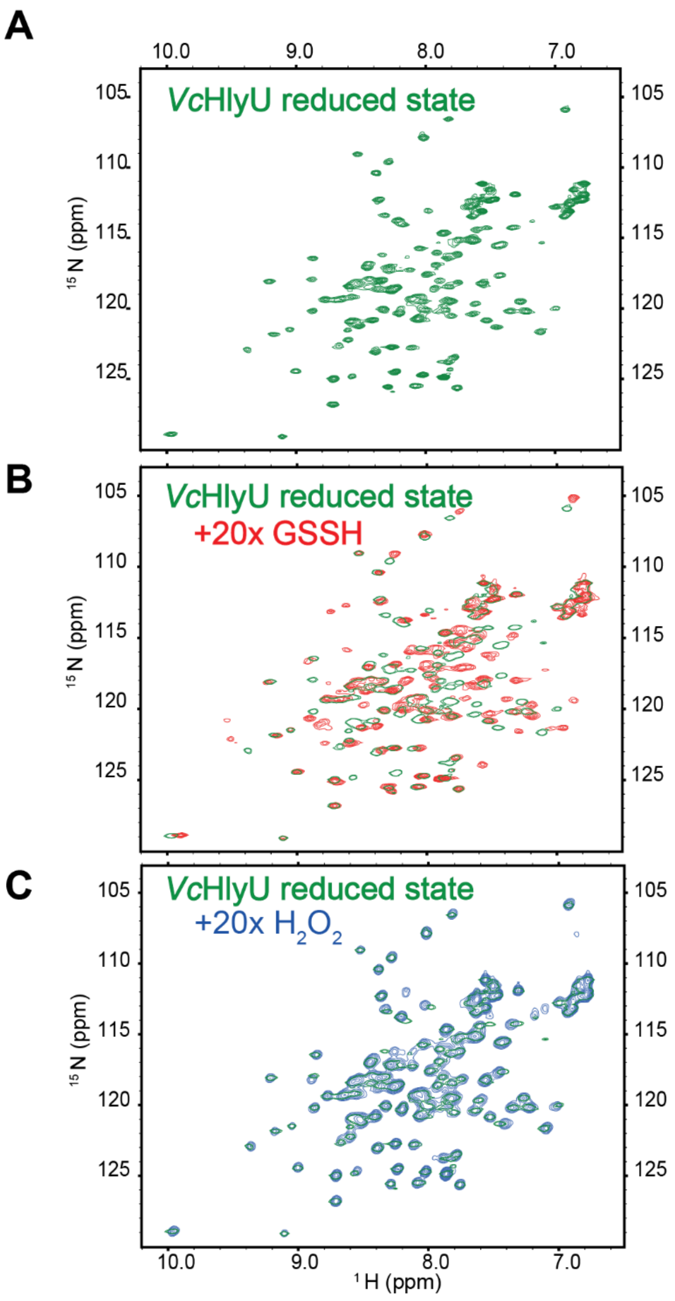


**Figure S4. (A)** $^1$H, $^{15}$N HSQC spectrum of reduced *Vibrio cholerae* HlyU. **(B)** Overlay of $^1$H, $^{15}$N HSQC spectrum of reduced *Vibrio cholerae* HlyU (*green*) and pre-treated with 20x GSSH (*red*). **(C)** Overlay of $^1$H, $^{15}$N HSQC spectrum of reduced *Vibrio cholerae* HlyU (*green*) and pre-treated with 20x $H_2O_2$ (*blue*). All spectra were measured at 30 ºC using a 20mM MES pH 6, 250mM NaCl, 1mM EDTA buffer, with the addition of 2mM TCEP in the case of the reduced state.

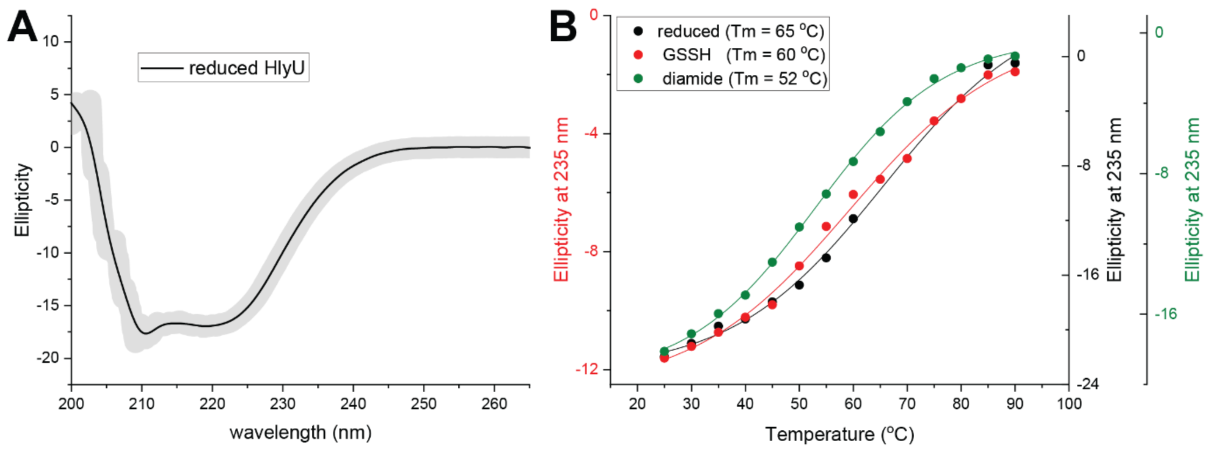


**Figure S5.** **(A)** Far-UV CD spectra of reduced HlyU. **(B)** Temperature-induced conformational transitions observed as changes in ellipticity at 232 nm in the 25–90 ºC temperature range for reduced (*black*), tetrasulfide (*red*) and diamide treated (disulfide, *green*) crosslinked HlyU. The line indicates a sigmoidal fitting used to obtain the melting temperature. All spectra were measured using in 25 mM HEPES, pH 7.0, 200 mM NaCl, 1 mM EDTA, with the addition of 1 mM TCEP in the case of the reduced state.

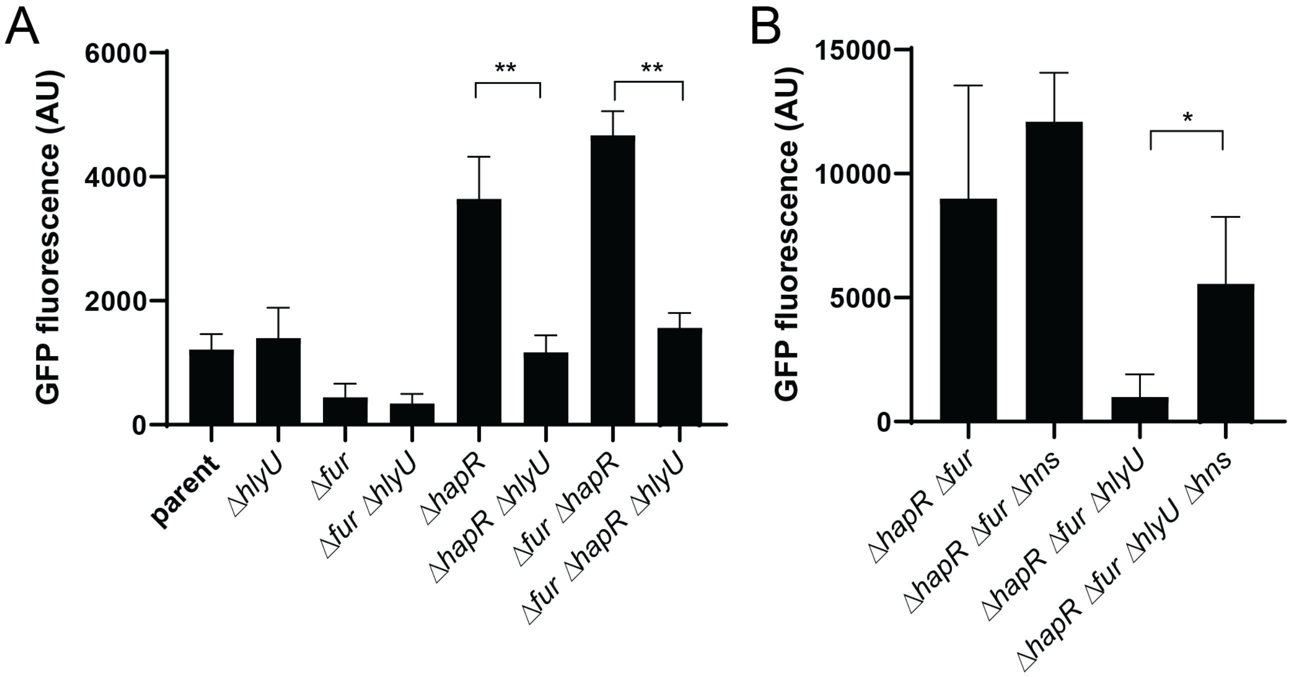


**Figure S6.** Assessing HlyU activity *in vivo* using a $P_{hlyA}$-GFP transcriptional reporter. Fluorescence of the indicated *V. cholerae* strains was measured to assess the impact of **(A)** HapR / Fur, and **(B)** HNS on HlyU-dependent activation of $P_{hlyA}$. Data are from four independent biological replicates and shown as the mean ± SD. Statistical significance was established using a unpaired parametric *t*-test (**p<0.01, *p<0.05).

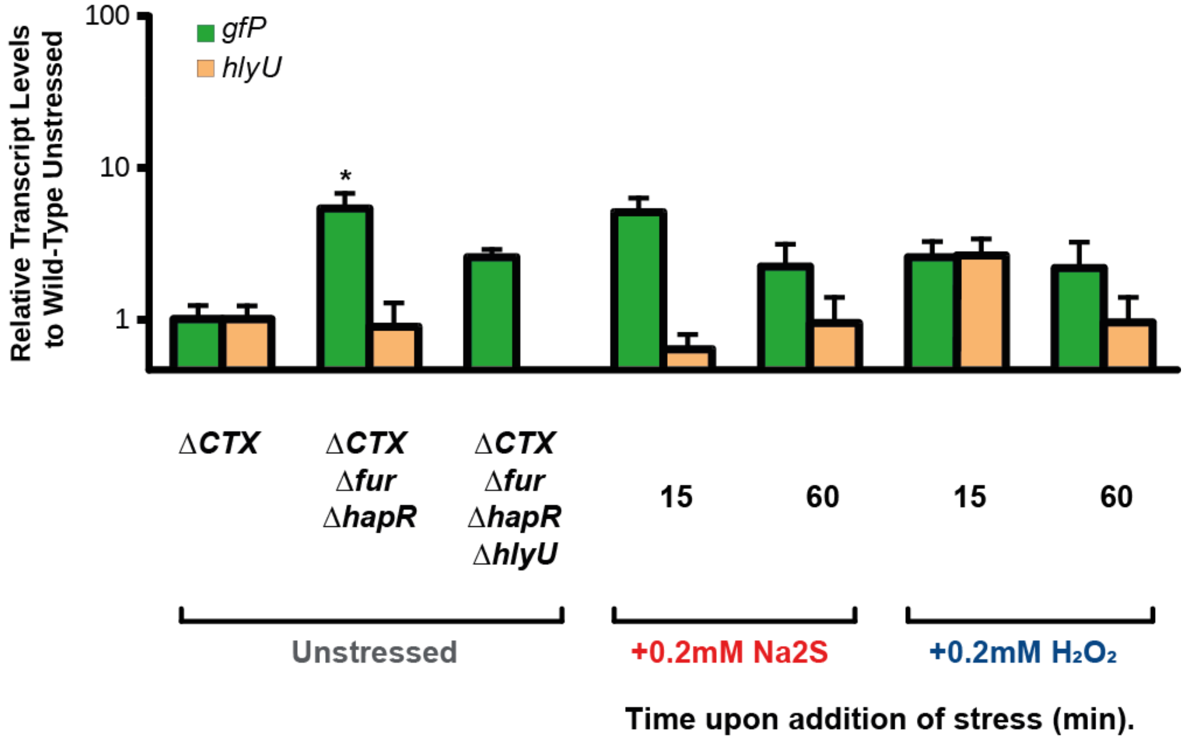


**Figure S7.** Fold changes transcripts levels of *Vc hlyU* and *gfP* followed by quantitative RT-PCR performed over a ΔCTX Δfur ΔhapR *V. cholerae* strain with the addition of $Na_2S$ or $H_2O_2$. Transcript values were normalized relative to the transcription level of *recA*. The values correspond to transcript levels relative to wild-type unstressed (WT UN) and are shown as mean ± SD from replicate cultures. Statistical significance was stablished using a paired t test relative to WT UN under the same conditions (**p<0.01, *p<0.05).

A

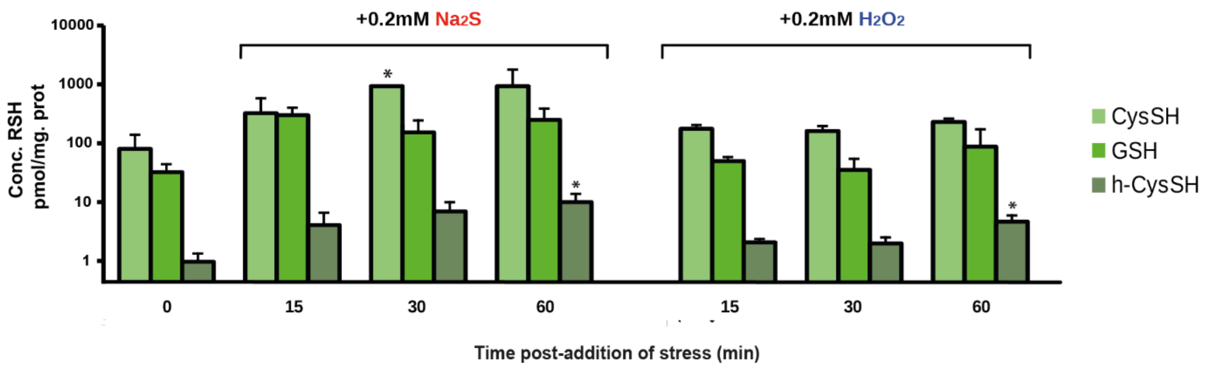


B

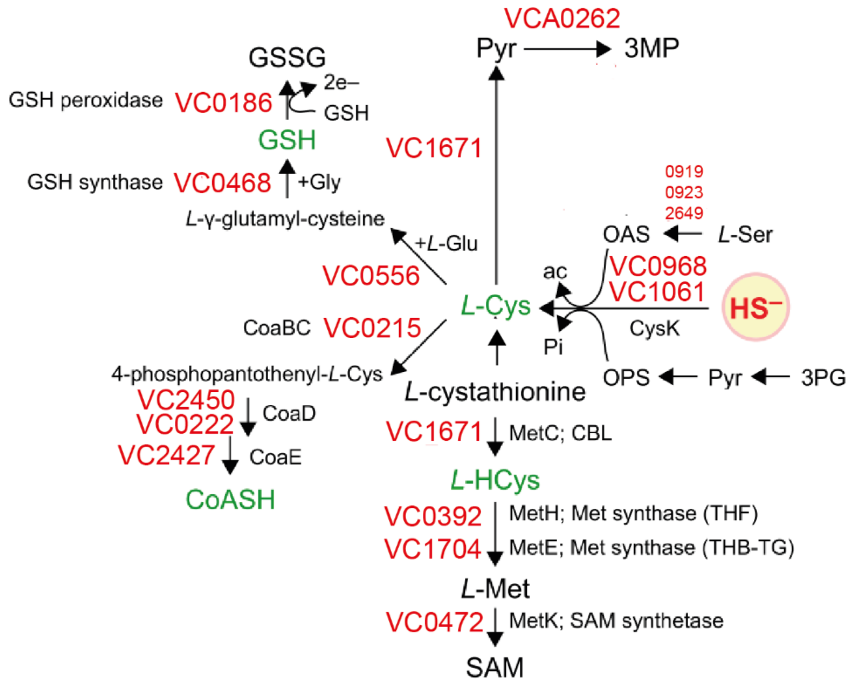


**Figure S8. (A)** Endogenous concentrations of LMW thiols before and after the addition of $Na_2S$ and $H_2O_2$ to mid-log-phase cultures (*, $P< 0.05$ using a paired t test relative to WT UN under the same conditions) determined using HPEIAM as capping agent. **(B)** *V. cholerae* genes encoding proteins associated with the biosynthesis of LMW thiols. OPS, O-phospho-L-serine; OAS, O-acetyl-L-serine; ac, acetate; pyr, pyruvate; CBL, cystathonine-γ-lyase. Adapted from reference(48).

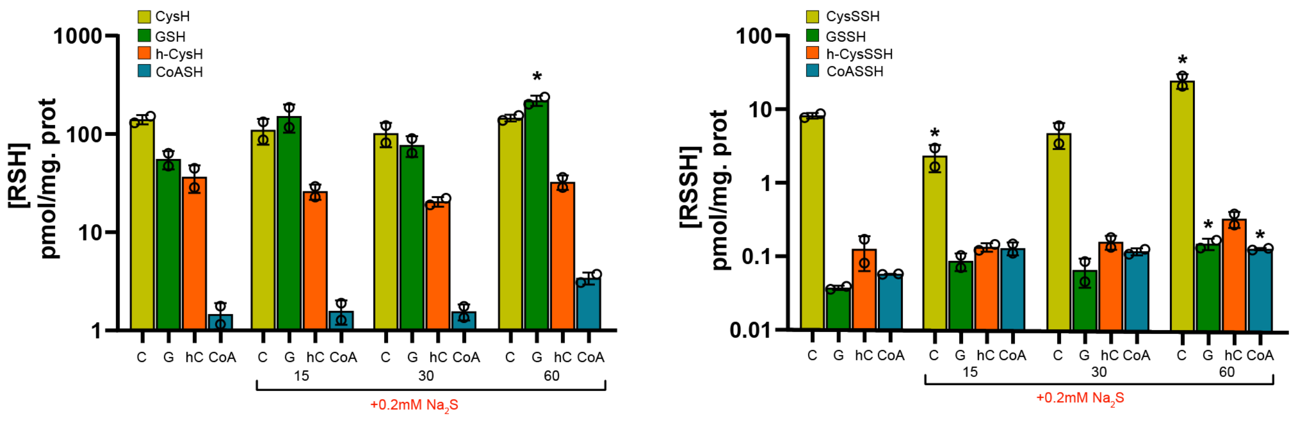


**Figure S9. (A)** Endogenous concentrations of LMW thiols before and after the addition of 0.2 mM $Na_2S$ to mid-log-phase cultures (*, $p < 0.05$) determined using mBBr as capping agent. **(B)** Endogenous concentrations of LMW persulfides before and after the addition of $Na_2S$ to mid-log-phase cultures (*, $p < 0.05$ using a paired t test relative to WT UN under the same conditions) determined using mBBr as capping agent.

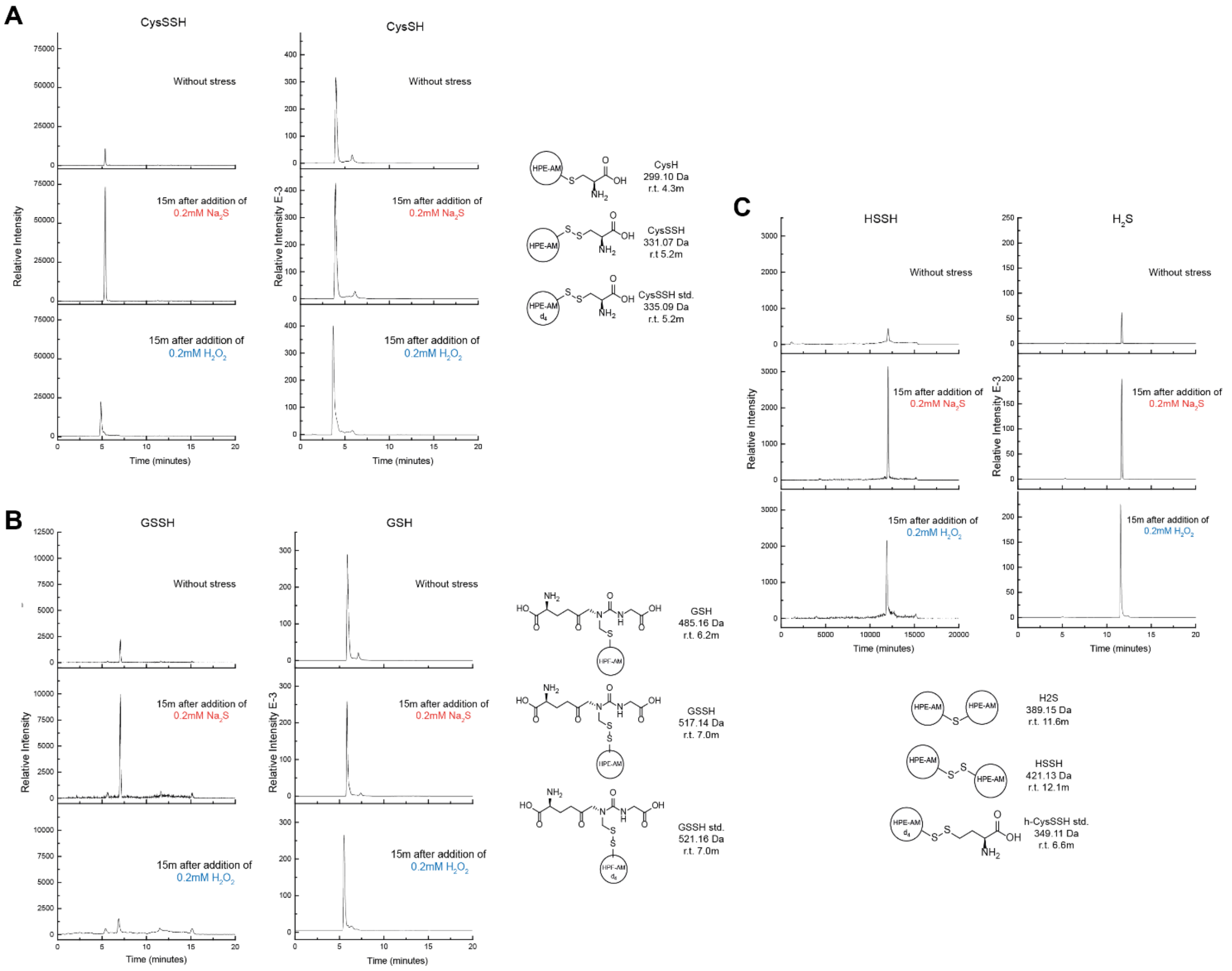


**Figure S10. (A)** Extracted ion chromatograms corresponding to cysteine thiol and cysteine persulfide (top panel) in unstressed cells and following 15 min of applied stress ($Na_2S$, middle panel and $H_2O_2$, bottom panel). **(B)** Extracted ion chromatograms corresponding to glutathione thiol and glutathione persulfide. **(C)** Extracted ion chromatograms corresponding to inorganic sulfide and disulfide. The peaks show in each case the relative intensity of the HPE-IAM labelled metabolites, relative to the intensity of the internal standard. On the right, we show the molecular structure, molecular weight and retention time of the metabolites analyzed in the panels, together with the internal standard used in each case.
